# SnRK2.4 and SnRK2.10 redundantly control developmental leaf senescence by sustaining ABA production and signaling

**DOI:** 10.1101/2025.03.25.645185

**Authors:** Anna Anielska-Mazur, Julia Rachowka, Lidia Polkowska-Kowalczyk, Michal Krzyszton, Dominika Cieslak, Radoslaw Mazur, Maria Bucholc, Jolanta Rakowska, Dominika Trzmiel, Paulina Stachula, Mateusz Olechowski, Szymon Swiezewski, Grazyna Dobrowolska, Anna Kulik

**Affiliations:** Institute of Biochemistry and Biophysics, Polish Academy of Sciences, Pawinskiego 5a, 02-106 Warsaw, Poland; University of Warsaw, Faculty of Biology, Institute of Biochemistry, Miecznikowa 1, 02-096 Warsaw, Poland; Centre of New Technologies, University of Warsaw, Banacha 2c, 02-097 Warsaw, Poland; University of Warsaw, Faculty of Biology, Institute of Genetics and Biotechnology, Pawinskiego 5a, Warsaw 02-106, Poland; Warsaw University of Life Sciences - SGGW, Nowoursynowska 166, 02-787 Warsaw, Poland

## Abstract

Plants constantly and precisely control their growth by inducing distinct developmental programs to survive and produce high-quality offspring in the changing environment. The fine-tuning of the development according to endogenous and environmental signals requires exact intercellular signaling and a balanced response. Kinases of the Sucrose non-fermenting-1-Related protein Kinases type 2 (SnRK2s) family primarily take part in the response and adaptation to environmental stress factors. Notably, here we show that two ABA-non-activated SnRK2s, SnRK2.4 and SnRK2.10, are also activated in non-stress conditions in developmentally senescing leaves of *Arabidopsis thaliana*. Phenotypic, biochemical, and molecular analyses performed on single *snrk2.4* or *snrk2.10*, and double *snrk2.4/2.10* kinase mutants showed that SnRK2.4 and SnRK2.10, acting redundantly, promote developmental leaf senescence. Further, SnRK2.4 and SnRK2.10 enhance ABA accumulation in senescing leaves by inducing *NCED2*, one of key ABA biosynthesis-related genes. The two kinases induce developmental leaf senescence by modulating the expression of multiple ABA-responsive, osmotic stress, and senescence-related genes, such as the senescence master regulators *ORE1*, *ORS1*, *WRKY33, WRKY75,* and *ANAC087.* Furthermore, we show that SnRK2.4 and SnRK2.10 act upstream of MAPK signaling by enhancing the expression and activity of MAPKKK18, a senescence-inducing kinase. These results document a new regulatory function of SnRK2.4 and SnRK2.10: they are activated in Arabidopsis leaves in response to endogenous signals and redundantly induce developmental leaf senescence by stimulating ABA production and sustaining major ABA-dependent and -independent signaling pathways.

## INTRODUCTION

All living organisms, including plants, undergo dynamic changes during their lifetime. Senescence is the last stage of development and leads to a controlled death of cells, tissues, organs and, in some cases, the whole organism. In nature, senescence is one of the critical processes conditioning plant fitness and survival. The gradual degradation and remobilization of cellular components from dying cells to new tissues significantly reduces energy loss and conserves vital macro- and micronutrients (Woo et al., 2013; Maillard et al., 2015; Kim et al., 2016). This strategy constitutes one of the fundamental survival mechanisms for both annual and perennial plants and their offspring during the changing seasons. Leaf senescence in particular is of major importance in the plant life cycle and therefore is also of agricultural interest (Munne-Bosh, 2008; Kim et al., 2016). Leaf senescence can be induced by age/development-dependent factors correlated with the organ’s lifetime (for review see Woo et al., 2013; Kim et al., 2016; Liebsh and Keech, 2016), by the plant entering the reproductive phase of growth (Woo et al., 2013), or by external factors such as water deficit and salt stress (Allu et al.,2014; Kim et al., 2016), light stress (Song et al., 2014; Kim et al., 2016; for review see Liebsch and Keech, 2016;), nutrient deficiency (for review see Maillard et al., 2015), pathogen attack (for review see Zhang et al., 2020), and others. The control of leaf senescence involves highly complex genetic programs fine-tuned by multiple layers of regulation, including chromatin remodeling, and transcriptional, post-transcriptional, translational, and post-translational regulation (Guo et al., 2021a). Leaf senescence is induced by phytohormones, mostly abscisic acid (ABA), ethylene, jasmonic acid, and salicylic acid (Zhao and Chan, 2016; for review see Guo et al., 2021a). Numerous transcription factors (TFs) are involved in this process, mainly those from large families: NAC (NAM/ATAF/CUC), WRKY, and MYB (for review see Kim et al., 2016; Guo et al., 2021a; Zhang et al., 2021; Ahmad et al., 2024). They control the expression of thousands of senescence-associated genes (SAGs) in a tightly orchestrated manner (Guo et al., 2004; for review see Kim et al., 2016). Phenotypic analyses using insertion mutants of selected genes encoding protein kinases and phosphatases have revealed that they are also involved in the regulation of leaf senescence (Yang et al., 2022). Among them, protein kinases belonging to distinct families were identified, e.g., Mitogen-Activated Protein Kinases (MAPKs) (Zhou et al., 2009; Matsuoka et al., 2015; Li et al., 2024), Histidine Kinases (Kim et al., 2006), Receptor and Receptor-like Protein Kinases (Lee et al., 2011; Xu et al., 2011; Zhang et al., 2024), Calcium Dependent Protein Kinases (CDPKs) (Durian et al., 2020), and selected Sucrose non-fermenting 1-Related protein Kinases (SnRK2s) (Gao et al., 2016). However, no protein kinases have been shown to undergo activation during developmental leaf senescence *per se*.

Among the many kinase types, the SnRKs have attracted particular interest in recent years. They are classified in the SNF1/AMPK family which includes among others, the SNF1 (Sucrose Non-Fermenting-1) kinase from the yeast *Saccharomyces cerevisiae* and mammalian AMPK (AMP-activated Protein Kinase). In plants, they are divided into three subfamilies: SnRK1, SnRK2, and SnRK3. SnRK2s are plant-specific Ser/Thr protein kinases with a molecular mass of about 40 kDa. They have been identified in *Arabidopsis thaliana* (Boudsocq et al., 2004) and every plant species tested so far (for review see Kulik et al., 2011; Yoshiaki al., 2021; Shinozawa et al., 2019). In Arabidopsis, the SnRK2 subfamily is further divided into three groups: group 1 (SnRK2.1/2.4/2.5/2.9/2.10) – kinases not activated by ABA, group 2 (SnRK2.7/2.8) – weakly activated by ABA, and group 3 (SnRK2.2/2.3/2.6) - kinases strongly activated by ABA (for review see Kulik et al., 2011). The best-characterized SnRK2s, those belonging to group 3 (for review see Kulik et al., 2011; Fàbregas et al., 2020), regulate the functioning of the SLAC1 and KAT1 ion channels (Lee et al., 2009; Sato et al., 2009) involved in stomatal closure (Mustilli et al., 2002) and control seed dormancy and germination (Fujii et al., 2007; Nakashima et al., 2009), expression of stress-related genes (Fujii et al., 2007), and miRNA biogenesis (Yan et al., 2017). Their involvement in regulating plant growth, development, and leaf senescence has also been suggested (Fuji and Zhu, 2009; Zheng et al., 2010; Yoshida et al., 2019). The physiological role of group 1 SnRK2s is the least understood. Besides salinity and osmotic stress, they are also activated by cadmium ions, the fungal elicitor cryptogein, nitric oxide, and reactive oxygen species (ROS) (Wawer et al., 2010; Kulik et al., 2012 and 2015, Rachowka et al., 2023). Under salinity SnRK2.4 and SnRK2.10 control primary root growth and lateral root formation, respectively (McLoughlin et al., 2012). Further, the ABA-non-activated SnRK2s phosphorylate varicose (VCS), an mRNA decapping activator, and thus regulate mRNA decay and accumulation of osmotic stress-dependent transcripts (Soma et al., 2017; Kawa et al., 2020), and the ERD14 dehydrin, which modulates its subcellular localization under salt stress (Maszkowska et al., 2019). Finally, SnRK2.4 and SnRK2.10 are involved in the maintenance of ROS homeostasis under salinity and cadmium ion stresses (Kulik et al., 2012; Szymanska et al., 2019; Mazur et al., 2021). However, an involvement of the ABA-non-activated SnRK2s in the regulation of plant development and growth under favorable growth conditions are scarce (Zheng et al., 2010; Fujii et al., 2011; Kawa et al., 2020).

## RESULTS

### SnRK2.4 and SnRK2.10 redundantly affect growth and development

To investigate a possible role(s) of the ABA-non-activated SnRK2s in the regulation of plant development in the absence of stress we followed the growth of rosette leaves of SnRK2.4 and SnRK2.10 (the two best-characterized representatives of group 1 SnRK2s) knock-out mutants. Since SnRK2.4 and SnRK2.10 have been found to be (at least partially) redundant in their well-established functions (Kawa et al., 2020), in addition to the two single mutants we also used the double *snrk2.4-1/snrk2.10-1* mutant (called here snrk2.4/2.10 mutant). Neither of the single mutants showed any difference in their growth and development compared with the wild type. In contrast, the double *snrk2.4/2.10* mutant differed from the wild type (and the two single mutants) in both its overall growth and some aspects of development.

Upon reaching full development, as defined by their flowering, the *snrk2.4/2.10* mutant was a significantly delayed in yellowing of its rosette leaves relative to that in the single *snrk2.4*, *snrk2.10* mutants and wt plants (Fig. 1A, B). Leaf yellowing is the first visible symptom of senescence due to compromised chloroplast functioning reflected by enhanced degradation of chlorophylls in a multistep catabolic pathway called PAO (Pheophorbide *a* oxygenase/phyllobilin pathway) (Pružinská et al., 2005; for review see Zhu et al., 2017). Another clearly visible characteristic of the *snrk2.4/2.10* plants was that they were markedly smaller than the wt or the single mutants (Fig. 1C) but took roughly the same time to flower as wt plants (ca. 21 days)(Fig. 1D). Since *snrk2.4/2.10* plants have been shown to germinate with the same vigor as the single *snrk2.4* or *snrk2.10* mutants and wt plants (McLoughlin et al., 2012) and formed rosette leaves and flowered at the same time as wt plants (Fig. 1D, E), the observed “stay-green” phenotype (Fig. 1 A, B) suggested a redundant control of developmental leaf senescence by the two kinases studied. Because only the double *snrk2.4/2.10* mutant possess the “stay-green” phenotype, in next experiments we examined this plant line in more detail in comparison with wt plants.

**Figure 1.**
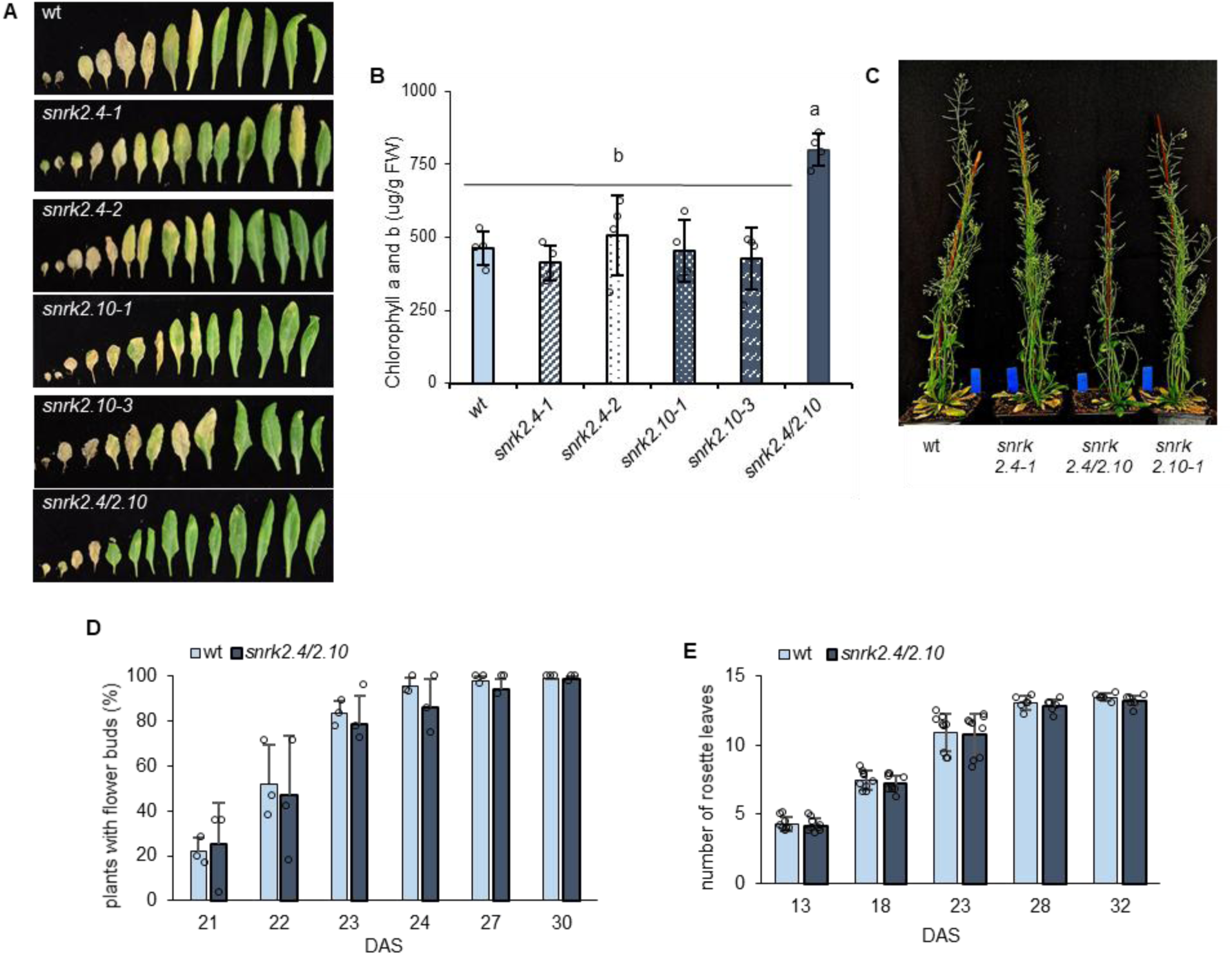
Impact of a deficit of SnRK2.4 and/or SnRK2.10 on Arabidopsis growth and development. A. Delayed leaf senescence in the *snrk2.4/2.10* mutant compared to *snrk2.4*, *snrk2.10,* and wt plants. Leaves detached from 45 days after sowing plants (45 DAS) are arranged in order from the oldest leaf (bottom) to the youngest one; cotyledons are not included. A representative photograph from three independent experiments with four biological replicates each is shown. B. Chlorophyll a and b content in 9th rosette leaves of wt, *snrk2.4*, *snrk2.10* and *snrk2.4/2.10* plants 45 DAS. A representative graph (mean ±SD) from three independent experiments with thirty plants of each line per experiment is shown. Different letters mark significantly different values as determined by ANOVA and Tukey post hoc test (p < 0.05). C. Morphology of wt, *snrk2.4-1*, *snrk2.10-1*, and *snrk2.4/2.10* 45 DAS plants cultivated in optimal conditions. A representative photograph from three independent experiments with five biological replicates each is shown. D. The rate of formation of the first flower bud in wild-type and *snrk2.4/2.10* plants. Data are represented as mean ±SD from three independent experiments with 30-50 plants per experiment. E. The rate of formation of successive rosette leaves in wild-type and snrk2.4/2.10 plants between 13 and 32 DAS. A representative graph (mean ±SD) from three independent experiments with 12 plants of each line per experiment is shown.

### SnRK2.4 and SnRK2.10 promote developmental leaf senescence

Before attempting to determine the mode of the involvement of SnRK2.4 and SnRK2.10 in the control of developmental leaf senescence we verified that the yellowing leaves do indeed undergo typical senescence. To this end we determined the level of expression of two well-characterized senescence marker genes *ORE1* (*ORESARA1*) and *SAG12* (*Senescence Associated Gene 12*) in the 6^th^ rosette leaves collected at different developmental stages: young, mature, early (up to 10% of the leaf blade yellowed), and late senescence. As expected, the *ORE1* and *SAG12* transcripts were hardly detectable in young and mature leaves, but their levels rose significantly in early senescent ones and increased even further in the late senescent leaves (Fig. 2A, B). In contrast, the expression of *SnRK2.4* and *SnRK2.10* increased markedly already with the leaf maturation and remained at a roughly similar high level throughout senescence, with some differences between the two genes (Fig. 2C, D).

**Figure 2.**
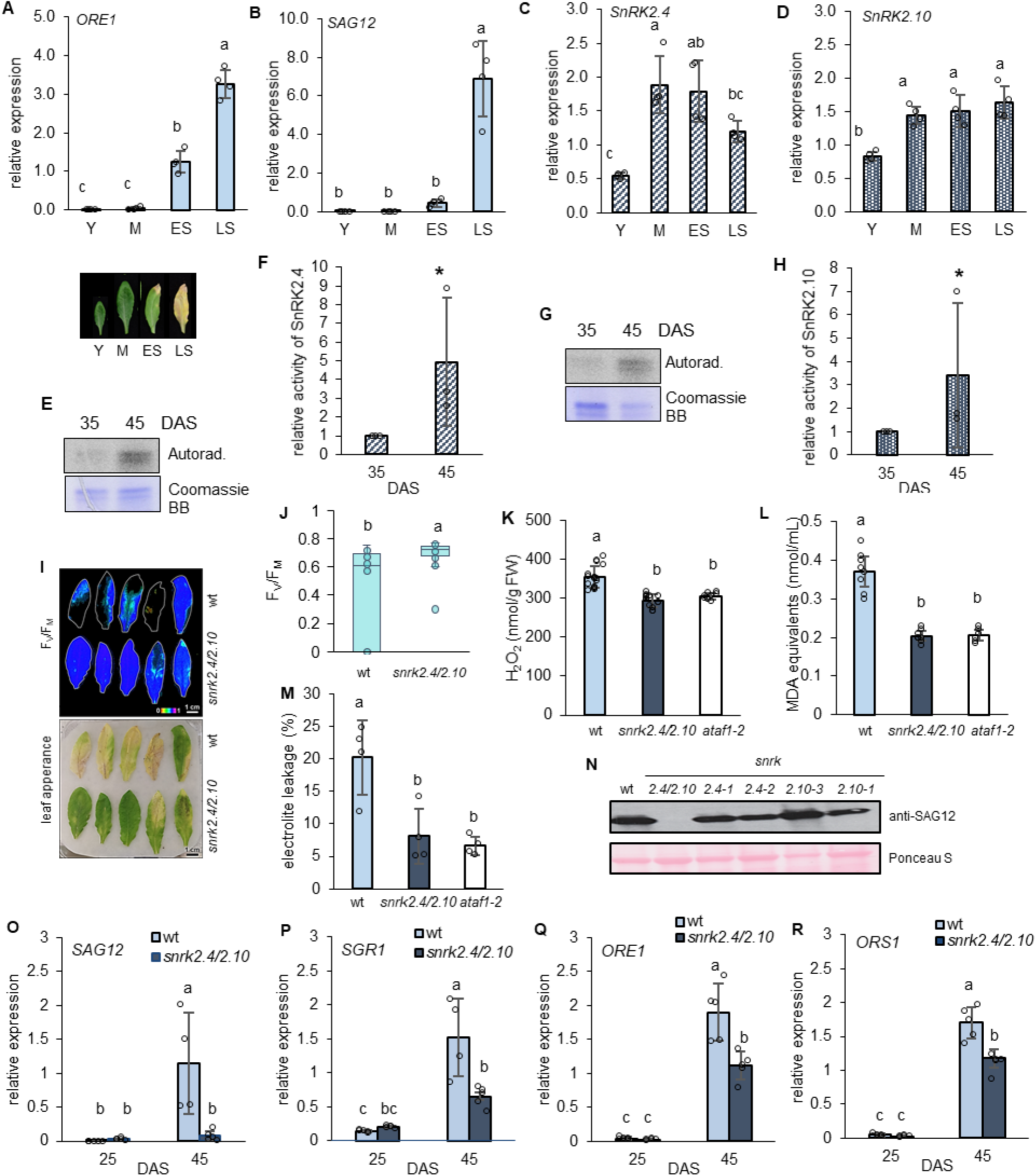
Developmental leaf senescence in *snrk2.4/2.10* mutant plants. A-D. Expression of *ORE1* (A), *SAG12* (B), *SnRK2.4* (C), and *SnRK2.10* (D) in the 6^th^ rosette leaves of wt plants collected at different developmental stages – young (Y), mature (M), early senescence (ES), and late senescence (LT). Representative transcripts were quantified by RT-qPCR relative to the *PEX4* housekeeping gene. Data are represented as mean ±SD from three independent experiments with ten plants per experiment. The representative image of leaves at different stages of development is presented E and G. Kinase activity of SnRK2.4 in *pSnRK2.4::SnRK2.4-YFP* (E) and of SnRK.10 in *pSnRK2.10::SnRK2.10-YFP* (G) transgenic plants. The activity was determined by immunocomplex kinase activity assay using anti-GFP antibodies and MBP as a substrate. 8^th^-10^th^ rosette leaves from plants 45 DAS were analyzed. Autorad. – autoradiogram. The experiment was repeated thrice with eight to ten plants per sample for each repetition. A representative image is shown. F and H. Quantification of the intensity of ^32^P-labeled MBP bands phosphorylated by SnRK2.4-YFP (shown in E) and by SnRK2.10-YFP (shown in G). Band intensity was determined using ImageJ and normalized against Coomassie Brilliant Blue (Coomassie BB) stained gel. The values shown are relative to wt plants 35 DAS. Means ±SD from three independent experiments are shown. I. Representative images of 9^th^ rosette leaves from wt and *snrk2.4/2.10* plants 45 DAS. Chlorophyll *a* fluorescence determination was performed using an Imaging-PAM chlorophyll fluorescence system. Upper panel shows the F_V_/F_M_ ratio and the lower one - leaf morphology. Three independent experiments were performed in triplicate with five leaves per sample. J. F_V_/F_M_ ratio in wt and *snrk2.4/2.10* plants 45 DAS. Chlorophyll *a* fluorescence determination was performed as in I. Mean values ±SD of three independent experiments, performed in triplicate with five leaves per sample, are shown. K. Hydrogen peroxide accumulation in 9^th^ rosette leaves of wt and *snrk2.4/2.10* plants 45 DAS. H_2_O_2_ was quantified using Amplex Red assay. Mean values ±SD of three or four independent experiments with eight to ten leaves per sample for each repetition, are shown. L. Malondialdehyde (MDA) accumulation in 9^th^ rosette leaves of wt and *snrk2.4/2.10* plants 45 DAS. Mean values ±SD of three independent experiments, performed in triplicate with eight to ten leaves per sample, are shown. M. Electrolyte leakage from 9^th^-10^th^ rosette leaves of wt, *snrk2.4/2.10,* and *ataf1-2* plants 45 DAS. Electrolyte leakage was determined by the conductivity measurement. Mean values ±SD of four independent experiments, performed in triplicate with five leaves per sample, are shown. N. Accumulation of SAG12 protein in 9^th^ rosette leaves of wt, *snrk2.4*, *snrk2.10*, and *snrk2.4/2.10* plants 45 DAS. SAG12 was determined in leaf extracts by western blotting with anti-SAG12 antibodies. A representative image of three independent repeats showing similar results is presented. O-R. Expression of senescence marker genes *SAG12* (O)*, SGR1* (P)*, ORE1* (Q), and *ORS1* (R) in 9^th^ rosette leaves of wt and *snrk2.4/2.10* plants 25 and 45 DAS. The transcripts were quantified by RT-qPCR relative to the *PEX4* housekeeping gene. Data are represented as mean ±SD from five independent experiments with ten plants per experiment. In all graphs, different letters mark significantly different values as determined by ANOVA and Tukey post hoc test (*p* < 0.05), except for F-H, where the asterisks indicate significant differences from the control (Student’s *t*-test; *p* < 0.05).

To determine the enzymatic activity of the two kinases upon leaf senescence we used transgenic plants expressing them as fusion proteins with YFP (*pSnRK2.4::SnRK2.4-YFP* or *pSnRK2.10::SnRK2.10-YFP*) in an appropriate insertion mutant background. We compared mature but non-senescent leaves (obtained from plants 35 days after sowing (DAS)) with senescent leaves showing both chlorophyll loss and high expression of the senescence marker genes (Fig. S1) from 45-day-old plants. In both cases, the 8^th^ to 10^th^ rosette leaves were taken for analysis. The fusion proteins were immunoprecipitated with anti-GFP antibodies and subjected to a kinase assay with MBP and [γ-^32^P]ATP as substrates followed by SDS-PAGE and quantification of radiolabeled MBP. The activity of both kinases was significantly elevated in the senescing leaves of plants 45 DAS compared with the non-senescing ones (Fig. 2E-H). The activation of SnRK2.4 and SnRK2.10 in senescing leaves indicates their involvement in regulating organ development in optimal growth conditions, in the absence of any exogenous stimuli. Since developmental leaf senescence is characterized by highly complex but not always specific tissue changes (Bresson et al., 2018), we analyzed several commonly used physiological, biochemical, and molecular markers to investigate the role of SnRK2.4 and SnRK2.10 in the regulation of this process.

One of the earliest changes in senescent leaves comprises inhibition of the expression of genes encoding proteins involved in photosynthesis with a simultaneous degradation of chloroplast proteins and chlorophylls. The chlorophyll degradation compromises the integrity of the photosynthetic apparatus and its ability to absorb light energy, which in turn leads to a photooxidative stress with a high production of ROS. The ROS cause an immediate drop of the photosynthetic yield and an increase in non-photochemical quenching (NPQ) (Woo et al., 2013; Pintó-Marijuan and Munné-Bosch, 2014). The F_V_/F_M_ parameter (reflecting the quantum efficiency of photosystem II in the dark) showed a mosaic distribution in a single senescing 9^th^ rosette leaf from both the *snrk2.4/snrk2.10* mutant and wt plants (Fig. 2I). However, the mean F_V_/F_M_ value in senescent leaves of the *snrk2.4/2.10* mutant was significantly higher than that in wild-type plants, indicating a higher overall photosynthetic efficiency and less damage to the photosynthetic apparatus in the mutant (Fig. 2J). In Arabidopsis, the initiation of developmental leaf senescence is accompanied by an increase in the intracellular H_2_O_2_ concentration caused by a reduced activity of catalases and APX1 (ascorbate peroxidase 1) and enhanced production of ROS in chloroplasts, peroxisomes, membranes, mitochondria, etc. (Allu et al., 2014; Pintó-Marijuan and Munné-Bosch, 2014; Jajic et al., 2015). As a consequence, membrane proteins and lipids are oxidized and degraded, which induces the accumulation of a large variety of oxidation products such as malondialdehyde (MDA). Finally, the plasma membrane becomes permeable and collapses, a clear indicator of which is an increased electrolyte leakage in senescing leaves (Bresson et al., 2018). The accumulation of H_2_O_2_, MDA content, and electrolyte leakage determined in 9^th^ rosette leaves of *snrk2.4/2.10* plants 45 DAS were all significantly lower than in wt plants and corresponded to the values obtained for the *ataf1-2* mutant (Fig. 2K-M). ATAF1 (*Arabidopsis thaliana* ACTIVATING FACTOR 1) is a TF belonging to the NAC (NO APICAL MERISTEM/ARABIDOPSIS ACTIVATION FACTOR1 [ATAF1-2]/CUP-SHAPED COTYLEDON) family, known to be a major leaf senescence-promoting regulator, whose knockout mutants exhibit a substantially delayed developmental leaf senescence (Garapati et al., 2015).

The massive protein degradation occurring in senescing leaves is carried out by diverse senescence-induced proteases. Among them is SAG12, a cysteine protease that is strongly induced upon developmental leaf senescence but not during stress-induced leaf senescence (Noh and Amasino, 1999). Thus, SAG12 can be used as a specific marker for monitoring age- and development-related changes in leaves. We observed an abundance of SAG12 in the 9^th^ rosette leaves of wt plants and the single *snrk2.4* and *snrk2.10* mutants at 45 DAS, but not in the *snrk2.4/2.10* leaves (Fig. 2N). Finally, expression of the senescence marker genes *SAG12, SGR1, ORE1*, and *ORS1* was induced in the senescing 9^th^ rosette leaves of wt and *snrk2.4/2.10* plants 45 DAS compared with 25 DAS leaves, but this induction was significantly lower in *snrk2.4/2.10* leaves compared with wt plants (Fig. 2O-R). These data indicate that SnRK2.4 and SnRK2.10 are activated and promote developmental leaf senescence in *A. thaliana*. Since the both single mutants, *snrk2.4* and *snrk2.10,* do not differ from the wt as concerns leaf senescence, and only the double mutant shows an effect, one must conclude that SnRK2.4 and SnRK2.10 can fully replace each other, i.e., they act redundantly.

### SnRK2.4 and SnRK2.10 control expression of multiple Senescence Associated Genes (SAGs) in senescing leaves

To understand better the genome-wide regulation of developmental leaf senescence and its dependence on SnRK2.4/2.10 signaling, we performed 3’ RNA-seq in the 9^th^ rosette leaves of wt and *snrk2.4/2.10* plants 25 and 45 DAS. As many as 670 showed significantly different expression between wt and the double mutant (differentially expressed genes, DEGs) in senescent leaves and only 113 in the non-senescent ones (Fig. 3A and Table S1). These results indicate a minimal role of the kinases studied in the modulation of gene expression in young and mature leaves but a substantial such role in senescing leaves. Notably, of the 670 genes affected by SnRK2.4/2.10 in senescing leaves only 13 showed such a dependence also in non-senescing leaves (Fig. 3B). Not only the number of DEGs was much lower in mature leaves compared with the senescing ones, but also the extent of the difference in their expression levels between wt and *snrk2.4/2.10* was markedly smaller (Fig. 3C). These data clearly indicate only a minor effect of the two SnRK2 kinases on gene expression in non-senescent leaves and a major one, both quantitatively and qualitatively, in senescent leaves.

**Figure 3.**
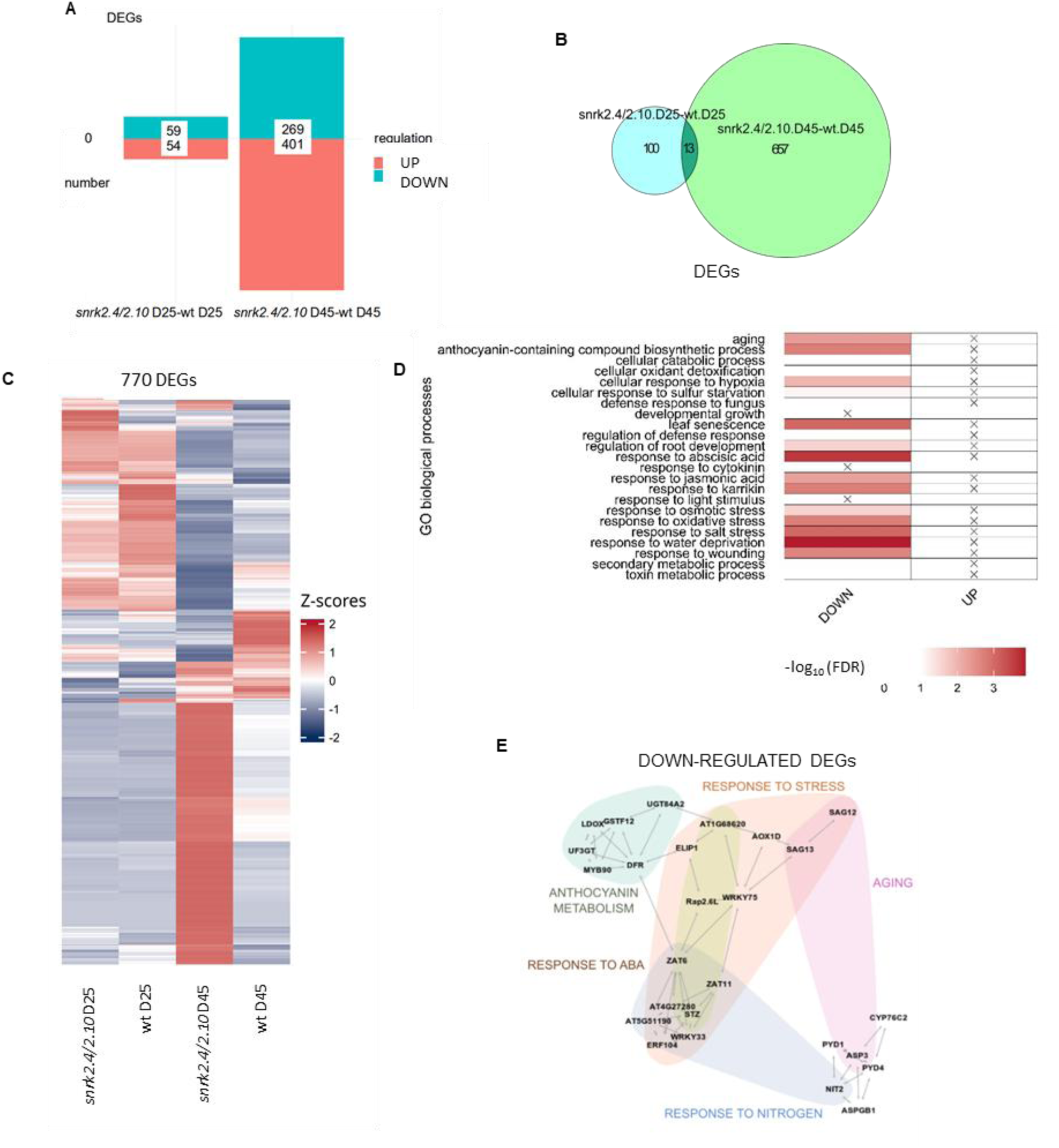
Impact of a deficit of SnRK2.4 and SnRK2.10 on expression of Senescence-Associated Genes. A. Up- and down-regulated DEGs in leaves of *snrk2.4/2.10* mutant *vs.* wt plants 25 and 45 DAS. B. Dependencies between DEGs in leaves of *snrk2.4/2.10 vs.* wt plants 25 and 45 DAS. Circle size reflects number of genes. C. Relative expression levels of DEGs in leaves of *snrk2.4/2.10* mutant *vs.* wt plants 25 and 45 DAS. The Z-score represents the deviation from the mean in standard deviation units. Red indicates a higher-than-mean expression level of all genes and blue – a lower one. D. Enriched GO terms from the “Biological process” among the DEGs in *snrk2.4/2.10* mutant *vs.* wt plants 25 and 45 DAS. Color intensity indicates the -log_10_ -transformed False Discovery Rate. All the terms shown were over-represented with a statistical significance of *p*<0.05. X mark an absence of DEGs for a given term. E. Down-regulated hub genes identified in the network analysis for *snrk2.4/2.10* 45 DAS *vs.* 25 DAS. Genes identified in wt 45 DAS *vs.* 25 DAS were subtracted from the analysis. Colors mark manually assigned groups, as labeled.

A functional analysis of the genes controlled by SnRK2.4/2.10 in senescing leaves revealed an overrepresentation of several gene ontology (GO) terms. These DEGs were mainly associated with response to water deprivation/osmotic stress, response to abscisic acid, leaf senescence, response to salt stress, response to wounding, response to oxidative stress, and anthocyanin biosynthesis (Fig. 3D, Fig. S2, Table 1 and Table S1). The large representation of genes related to the responses to osmotic stress, oxidative stress, and salinity is consistent with the previously recognized role of SnRK2.4 and SnRK2.10 in conditioning plant resistance to water deficit and ROS homeostasis (McLoughlin et al., 2012; Kulik et al., 2012; Szymańska et al., 2019; Mazur et al., 2021; Rachowka et al., 2023). Similarly, the presence of overrepresented DEGs related to ‘leaf senescence’ and ‘aging’ is consistent with the previously published transcriptomic data from leaf development studies (Guo et al., 2004; Garapati et al., 2015; Woo et al., 2016). In contrast, the finding of a rather large group of DEGs (61 genes) involved in the response to abscisic acid is highly unexpected and could indicate that the studied kinases (which are not activated by ABA) could play a previously unknown role(s) in ABA sensing, signal transduction, or metabolism.

**Table 1.** List of 25 most down-regulated DEGs identified in leaves of *snrk2.4/2.10* mutant *vs* wt plants 45 DAS.

| gene | name | annotation | involved in | Log<sub>2</sub>FC | adj.pval |
| --- | --- | --- | --- | --- | --- |
| AT3G57510 | ADPG1 | Pectin lyase-like superfamily protein | anther dehiscence, cell wall modification involved in abscission, fruit dehiscence | -5.48 | 9.7E-04 |
| AT1G47510 | 5PTASE11 | Posphatidylinositol polyphosphate 5-phosphatase | phosphatidylinositol dephosphorylation, response to ABA, response to auxin, response to JA, response to salt stress | -5.22 | 5.55E-06 |
| AT1G10940 | SNRK2.4 | Serine/threonine-protein kinase SnRK2.4 | primary root development, response to osmotic stress, response to salt stress | -5.02 | 1.12E-08 |
| AT2G41850 | ADPG2 | Pectin lyase-like superfamily protein | anther dehiscence, cell wall modification involved in abscission, floral organ abscission, fruit dehiscence | -4.91 | 8.40E-03 |
| AT4G16740 | TPS03 | Terpene synthase | diterpenoid biosynthetic process, response to herbivore, response to jasmonic acid, response to wounding | -4.54 | 1.82E-04 |
| AT5G55410 | - | Lipid-transfer protein. Seed storage 2S albumin superfamily protein | systemic acquired resistance | -4.51 | 8.71E-04 |
| AT1G16950 | - | Transmembrane, extracellular signaling peptide | cellular response to nitrogen starvation, nitrate import, regulation of root development, response to ammonium ion, response to gibberellin, response to osmotic stress | -4.48 | 1.76E-06 |
| AT3G52780 | PAP20 | Purple acid phosphatases superfamily protein | unknown | -4.46 | 2.22E-03 |
| AT3G60140 | DIN2, SRG2 | Glycosyl hydrolase superfamily protein | carbohydrate metabolic process, glucosinolate catabolic process, response to salt stress, response to darkness, response to sugar and sulfur starvation | -4.38 | 3.66E-02 |
| AT2G14620 | XTH10 | Xyloglucan endotransglucosylase/hydrolase protein 10 | cell wall biogenesis, xyloglucan metabolic process | -4.29 | 01.18E-02 |
| AT1G11190 | BFN1 | Bifunctional nuclease that acts on both RNA and DNA | DNA catabolic process, floral organ senescence, leaf senescence | -4.27 | 5.98E-02 |
| AT1G68450 | PDE337 | VQ motif-containing protein | may be involved in chloroplast development | -4.22 | 7.22E-03 |
| AT1G28130 | GH3.17 | Indole-3-acetic acid-amido synthetase GH3.17 | intracellular auxin homeostasis, response to auxin | -4.21 | 6.26E-04 |
| AT3G03530 | NPC4 | Non-specific phospholipase C (PLC) | phospholipid catabolic process, response to sulfur deficiency, response to salt stress | -4.16 | 8.31E-05 |
| AT5G49600 | SVB6 | ABA-responsive SVB family gene | response to ABA | -3.96 | 2.32E-05 |
| AT3G14770 | SWEET2 | Bidirectional sugar transporter SWEET2 | carbohydrate transport, response to microbiota | -3.90 | 9.42E-04 |
| AT3G05400 | - | Major facilitator superfamily protein; sugar transporter | sugar transmembrane transport | -3.85 | 5.85E-04 |
| AT3G21500 | DXPS1 | Protein postulated to have 1-deoxy-D-xylulose 5-phosphate synthase activity | 1-deoxy-D-xylulose 5-phosphate biosynthetic process, chlorophyll biosynthetic process, terpenoid biosynthetic process | -3.84 | 6.95E-06 |
| AT1G58340 | ZF14 | Multidrug and toxin extrusion transporter | iron ion transport, regulation of plant organ formation, response to nematode, response to ABA, response to the absence of light, response to heat, response to osmotic stress, senescence | -3.72 | 1.06E-04 |
| AT5G24240 | ATPI4KGA MMA3, MOP9.5 | Phosphatidylinositol 3- and 4-kinase | regulation of flower development, response to abscisic acid, response to salt | -3.72 | 3.35E-04 |
| AT1G02310 | MAN1 | Glycosyl hydrolase superfamily protein | mano-oligosaccharide hydrolysis | -3.71 | 3.24E-04 |
| AT1G71520 | ERF020 | Ethylene-responsive transcription factor ERF020 | unknown | -3.67 | 1.29E-04 |
| AT4G18425 | DMP4 | transmembrane protein, putative (DUF679) | abscission, endomembrane system organization, plant organ senescence | -3.65 | 6.61E-03 |
| AT1G07610 | MT1C | Metallothionein-like protein 1C | response to copper ion, response to cadmium | -3.59 | 1.162E-03 |
| AT2G19500 | CKX2 | Cytokinin dehydrogenase 2 | cytokinin catabolic process, involved in organ development | -3.58 | 6.97E-03 |

By analyzing the genes specifically regulated by SnRK2.4 and SnRK2.10 in senescent leaves, we also identified hub genes and determined their connection network (Fig. 3E and Fig. S3). Among the hub genes down-regulated in the *snrk2.4/2.10* mutant relative to wt, genes encoding transcription factors such as *WRKY75*, *WRKY33*, *ZAT6*, *ZAT10 (STZ),* and *ZAT11* deserve special attention. *WRKY33* and *ZAT10* have already been recognized as hub genes controlling the response of plants to biotic and abiotic stress factors (Choura et al., 2015; Birkenbihl et al., 2017; Biniaz et al., 2022; Arjmand et al., 2023). Notably, *WRKY33* and *WRKY75* were previously found by us to be controlled by SnRK2.10 kinase in response to salt and oxidative stress (Rachowka et al., 2023). In agreement with the fact that developmental leaf senescence is regulated by multiple cross-talking signaling pathways and transcription factors, we also identified other SnRK2.4/2.10-dependent modules of senescence induction. For instance, upon leaf senescence SnRK2.4 and −2.10 were required for the induced expression of the senescence central regulator *ORE1* and its target genes such as *BFN1* (bifunctional nuclease 1), *SWEET15/SAG29*, *SGR1/NYE1*, *PAP20* (purple acid phosphatase), *At1g74010,* and others (Fig. 2, Table 1 and S1) (Balazadeh et al., 2010; Mattallana-Ramirez et al., 2013; Qiu et al., 2015). As previously shown, SnRK2.2, −2.3, and −2.6 can also induce ABA-dependent leaf senescence by up-regulating *ORE1* expression in an ABFs- and RAV1-dependent manner (Zhu et al., 2016). Further, the induction of expression of *ANAC087* and its direct targets *BFN1, NYE1,* and *CEP1* (cysteine endopeptidase 1) (Chen et al., 2023) was also reduced in senescing *snrk2.4/2.10* leaves. All these observations indicate that SnRK2.4 and SnRK2.10 promote leaf senescence and play a key role at the interface of the responses to a wide range of biotic and abiotic stress factors and internal stimuli by controlling hub stress-responsive and senescence-inducing genes such as *WRKY33, WRKY75, ORE1, ANAC087,* and others.

### SnRK2.4 and SnRK2.10 induce ABA production in senescing leaves

Abscisic acid is one of the major phytohormones involved, e.g., in seed development, root growth, stomatal aperture, and responses to water stress (Chen et al., 2020). ABA content increases during leaf senescence and exogenous application of ABA induces leaf senescence, and expression of multiple SAGs (Guo et al., 2021a). Since we observed a reduced expression of numerous ABA-responsive genes in senescing leaves in *snrk2.4/2.10* plants compared with the wild type, we reasoned that this effect could be caused by a reduced level of this phytohormone. Indeed, while the ABA content in the 9^th^ rosette leaves from the two plant lines was equally low at 23 and 36 DAS, at day 45 it significantly increased in wt leaves but remained unchanged in the *snrk2.4/2.10* mutant (Fig. 4A). The ABA biosynthesis takes place in chloroplasts and the cytosol by oxidative cleavage of the 11,12 double bond of 9-cis epoxy carotenoids (neoxanthin and/or violaxanthin) (Tan et al., 2003). Genes encoding the enzymes engaged in ABA biosynthesis are specifically expressed in different tissues in a conditions/stress-dependent manner (Tan et al., 2003). Among them, NCEDs (NINE-CIS-EPOXYCAROTENOID DIOXYGENASES) are the key rate-limiting enzymes in ABA biosynthesis (Thompson et al., 2000). During leaf senescence mainly *NCED2* and *NCED3* are induced (Xu et al., 2020). To decipher the mechanism underlying the lack of an induction of ABA accumulation in senescing leaves of the *snrk2.4/2.10* line we examined the expression of several genes involved in ABA synthesis and degradation. The induction of expression of *NCED2* and, to a lesser extent, *NCED3* and *AAO3*, was significantly lower in senescing leaves of the *snrk2.4/2.10* mutant compared with wild-type plants (Fig. 4B-C, Fig. S4C), while expression of *NCED4* was 3 times higher in the mutant compared with wt (Fig. 4D). The other genes tested were expressed at similar levels in the two lines (Fig. 4. This data reveals that SnRK2.4 and SnRK2.10 promote ABA production in developmentally senescing leaves by stimulating the expression of ABA-biosynthetic genes. However, SnRK2.4 and SnRK2.10 themselves are not activated by abscisic acid (Boudsocq et al., 2004). To check whether the two kinases could nevertheless be somehow involved in ABA-induced leaf senescence, we compared the responses of wt and *snrk2.4/2.10* leaves to ABA treatment.

**Figure 4.**
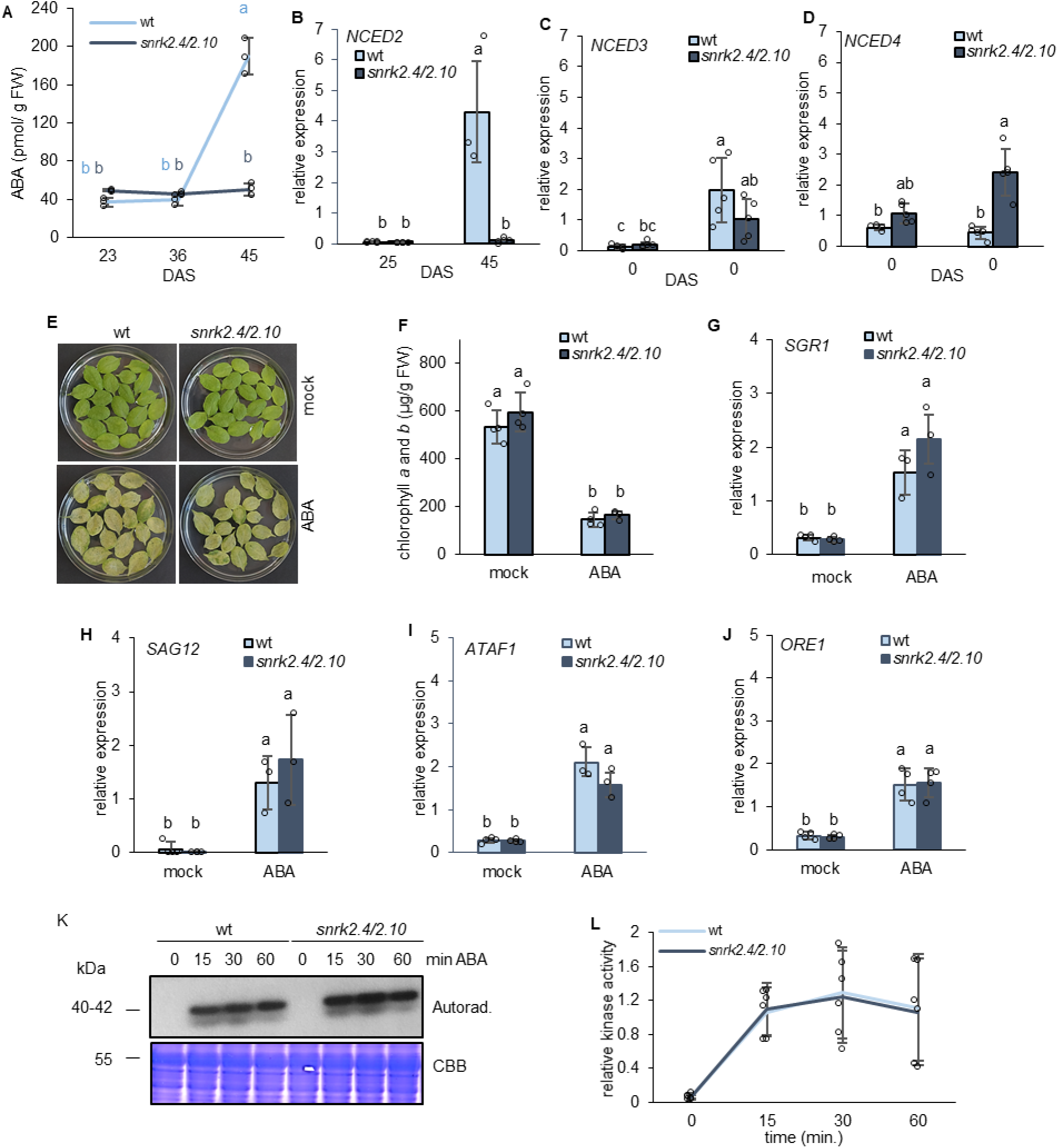
Impact of a deficit of SnRK2.4 and SnRK2.10 on ABA production and response of leaves to exogenous ABA. A. Accumulation of endogenous ABA in 9^th^ rosette leaves of wt and *snrk2.4/2.10* plants 23, 35, and 45 DAS. ABA content was determined by enzyme immunoassay. A representative graph (mean ±SD) from three independent experiments performed in triplicate, with eight-ten leaves per sample, is shown. Different letters mark significantly different values as determined by ANOVA and Tukey post hoc test (*p* < 0.05). B-D. Expression of *NCED2* (B), *NCED3* (C), and *NCED4* (D) in 9^th^ rosette leaves of wt and *snrk2.4/2.10* plants 25 and 45 DAS. The transcripts were quantified by RT-qPCR relative to *PEX4* housekeeping gene. Mean values ±SD of three to five independent experiments, with ten leaves per experiment, are shown. Different letters mark significantly different values as determined by ANOVA and Tukey post hoc test (*p* < 0.05). E. Phenotypes of wt and *snrk2.4/2.10* leaves after ABA treatment. Detached 3^rd^ and 4^th^ rosette leaves from plants 28 DAS were treated or not with 50 μM ABA for 2 days under continuous dim light. Representative photographs from three independent experiments performed in four replicates, with 20 leaves per sample, are shown. F. Chlorophyll *a* and *b* content in leaves of wt and *snrk2.4/2.10* plants presented in panel E. A representative graph (mean ±SD) from three independent experiments performed in four replicates, with 20 leaves per sample, is shown. Different letters mark significantly different values as determined by ANOVA and Tukey post hoc test (*p* < 0.05). G-J. Expression of senescence marker genes *SGR1* (G)*, SAG12* (H)*, ATAF1* (I), and *ORE1* (J) in leaves of wt and *snrk2.4/2.10* plants presented in panel E. The transcripts were quantified by RT-qPCR relative to the *PEX4* housekeeping gene. A representative graph (mean ±SD) from three independent experiments performed in four replicates, with 20 leaves per sample, is shown. Different letters mark significantly different values as determined by ANOVA and Tukey post hoc test (*p* < 0.05). K. Activity of ABA-induced kinases in wt and *snrk2.4/2.10* seedlings. Hydroponically grown ten-day-old seedlings were treated with 100 µM ABA for 0 to 60 min and protein extracts from whole seedlings were used for in-gel kinase activity assay using ABF2 fragment as a substrate. Representative photographs from three independent experiments with 100 seedlings per experiment, are shown. Autorad. – autoradiogram. CBB – Coomassie Brilliant Blue. L. Quantification of ^32^P-labeled ABF2 bands phosphorylated by ABA-activated kinases (shown in K). Band intensity was determined using ImageJ software and normalized against CBB-stained gel. Mean values ±SD of three independent experiments are shown.

The treatment caused a similar chlorophyll loss in both plant lines (Fig. 4E-F, Fig. S5). Also the expression of genes encoding enzymes responsible for chlorophyll (*SGR1*) and protein (*SAG12*) degradation and senescence-promoting TFs (*ATAF1* and *ORE1*) was induced similarly in wt and mutant leaves (Fig. 4G-J). Finally, we determined whether SnRK2.4 and SnRK2.10 affected the activity of ABA-activated SnRK2s (SnRK2.2, SnRK2.3, and SnRK2.6). The SnRK2.2, SnRK2.3, and SnRK2.6 were activated similarly in wt and *snrk2.4/2.10* mutant plants following ABA application and remained equally active throughout the experiment (Fig. 4K-L). Taken together, the results presented in this section show that SnRK2.4 and SnRK2.10 promote developmental leaf senescence by stimulating ABA production but they do not participate in the regulation of ABA-activated SnRK2s nor affect the leaf senescence caused by exogenous application of ABA.

### SnRK2.4 and SnRK2.10 impact MAPKKK18 signaling

Numerous protein kinases belonging to distinct families, including MAPK (Mitogen-Activated Protein Kinases), have been shown to contribute to leaf senescence (for review see Yang et al., 2022). To determine whether SnRK2.4 and SnRK2.10 play any role in regulating the functioning of the MAPK signaling pathways activated during leaf senescence, we determined the activity of selected kinases in 9^th^ rosette leaves of wild-type and *snrk2.4/2.10* plants. Having found that SnRK2.4 and SnRK2.10 aid ABA accumulation in senescing leaves, we focused our attention on the ABA-activated MAPK cascade, comprised of MAP3K17/18, MKK3, and four group C MAPKs, MPK1/2/7/14, which is atypically activated by the PYR/PYL/RCAR-SnRK2-PP2C ABA core signaling module through *de novo* synthesis of the MAP3Ks (Danquah et al., 2015). We selected for the analysis mitogen-activated protein kinase kinase kinase 18 (MAPKKK18) which is induced by ABA and is involved in drought stress resistance and developmental leaf senescence (Matsuoka et al., 2015; Mitula et al., 2015; Li et al., 2017; Zhao et al., 2023). Overexpression of MAPKKK18 has been shown to accelerate developmental and ABA-induced leaf senescence *via* its protein kinase activity (Matsuoka et al., 2015).

To determine whether SnRK2.4 and SnRK2.10 affect MAPKKK18 functioning during developmental leaf senescence, we assayed its expression and activity in non-senescent (35 DAS) and senescent (45 DAS) leaves of wild-type and *snrk2.4/2.10* plants. A basal activity of MAPKKK18 was found in leaves of both plant lines 35 DAS, while in senescing leaves, this activity was significantly induced in wt plants and remained unchanged in the *snrk2.4/2.10* mutant (Fig. 5A-B). Similarly, *MAPKKK18* expression was induced only in senescing leaves of wt plants, remaining at a low level in *snrk2.4/2.10* (Fig. 5C). This data indicates that the MAPKKK18-dependent signaling cascade is activated *in vivo* and acts downstream of SnRK2.4/2.10 in senescing leaves. To extend this observation the developmental leaf senescence was investigated in the 9^th^ rosette leaves of wild-type, *snrk2.4/2.10*, and *mapkkk18-1* mutant plants 45 DAS. Chlorophyll *a* and *b* levels were similar in the wt and *mapkkk18-1* lines and lower than in *snrk2.4/2.10* (Fig. 5D-E). In contrast, SAG12 protein accumulation was the highest in senescing wt leaves, while in the *snrk2.4/2.10* and *mapkkk18-1* mutants SAG12 was, respectively, absent or present in trace amounts only (Fig. 5F). This data suggests that the progression of developmental leaf senescence is more strongly affected by the SnRK2.4/2.10 kinases than by the MAPKKK18 cascade.

**Figure 5.**
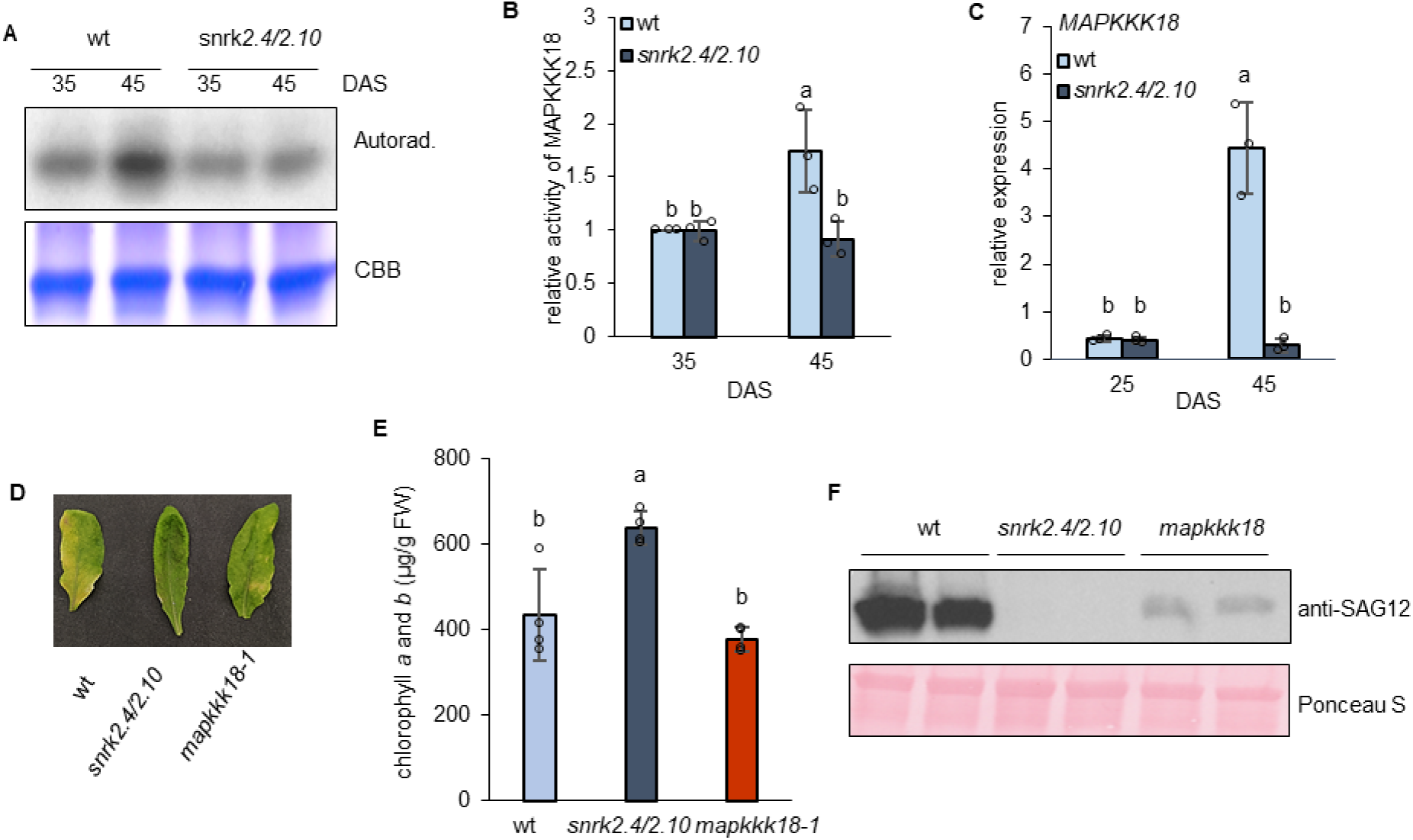
Impact of a deficit of SnRK2.4 and SnRK2.10 on MAPKKK18 signaling in senescing leaves. A. Activity of MAPKKK18 kinase in 8^th^-10th^th^ rosette leaves of wt and *snrk2.4/2.10* plants 35 and 45 DAS. Kinase activity was determined by immunocomplex kinase activity assay using anti-MAPKKK18 antibodies and MBP as a substrate. Representative photographs from three independent experiments, with eight plants per experiment, are shown. Autorad. – autoradiogram. CBB – Coomassie Brilliant Blue. B. Quantification of ^32^P-labeled MBP bands phosphorylated by MAPKKK18 (shown in A). Band intensity was determined using ImageJ software and normalized against CBB-stained gel. The values shown are relative to wt plants 35 DAS. Means ±SD from three independent experiments are shown. Different letters mark significantly different values as determined by ANOVA and Tukey post hoc test (*p* < 0.05). C. Expression of *MAPKKK18* in 9^th^ rosette leaves of wt and *snrk2.4/2.10* plants 25 and 45 DAS. *MAPKKK18* transcript was quantified by RT-qPCR relative to the *PEX4* housekeeping gene. Mean values ±SD of three independent experiments, with 15 leaves per experiment, are shown. Different letters mark significantly different values as determined by ANOVA and Tukey post hoc test (*p* < 0.05). D. Comparison of developmental leaf senescence in wt, *snrk2.4/2.10,* and *mapkkk18* mutants. Leaves detached from plants 45 DAS are arranged in order from the oldest leaf (bottom) to the youngest one; cotyledons are not included. A representative photograph from five independent biological replicates is shown. E. Chlorophyll *a* and *b* content in 9^th^ rosette leaves of wt, *snrk2.4/2.10,* and *mapkkk18* plants 45 DAS. Mean values ±SD of three independent experiments, with ten leaves per experiment, are shown. Different letters mark significantly different values as determined by ANOVA and Tukey post hoc test (*p* < 0.05). F. Accumulation of SAG12 protein in 9^th^ rosette leaves of wt, *snrk2.4*, *snrk2.10*, and *snrk2.4/2.10* plants 45 DAS. SAG12 was detected in leaf extracts by western blotting with anti-SAG12 antibodies. A representative image of five independent experiments, with ten leaves per experiment is presented.

Finally, we determined the activity of the ABA-activated kinases SnRK2.2/2.3/2.6 in senescing leaves of wt and *snrk2.4/2.10*. They were activated *in vivo* in the both lines (Fig. S6 A-B), albeit to a slightly lower extent in *snrk2.4/2.10,* suggesting that SnRK2.4 and SnRK2.10 could act upstream of SnRK2.2, −2.3 and −2.6.

## DISCUSSION

Leaf senescence leads to irreversible changes in cells eventually leading to their death, therefore it has to be tightly regulated (Guo et al., 2021a). The protein kinase signaling networks involved in the control of leaf senescence and their cross-talk *in vivo* are still poorly understood (Yang et al., 2022). Here, we showed that ABA-non-activated SnRK2s, SnRK2.4 and SnRK2.10, play a redundant role in inducing developmental leaf senescence in *Arabidopsis thaliana*, which finding increases the complexity of the relevant regulatory network even further. Leaves collected from a double *snrk2.4/2.10* knock-out mutant plants at 45 DAS presented a distinct stay-green phenotype when compared with wt leaves or the single *snrk2.4* or *snrk2.10* mutants, which all showed clear-cut signs of senescence, i.e., extensive yellowing. This difference was most likely caused by an impaired induction of the transcription factor *ORE1* up-regulating the chlorophyll degradation-related gene *SGR1* (*NYE1*)(Stay-Green 1/Non-Yellowing 1) (Qiu et al., 2015). Under senescence, the *SGR1/NYE1* transcript accumulates and the encoded protein acts as a key positive regulator of chlorophyll breakdown through its strong binding to both Light Harvesting Complex II (LHCII) and a set of six chlorophyll catabolic enzymes - CCEs (NYC1, NOL, HCAR, PPH, PAO, and RCCR)(Zhu et al., 2017). A series of biochemical, physiological, and molecular markers also indicated a delayed developmental leaf senescence in *snrk2.4/2.10* leaves. This observation correlated with the *in vivo* activation of the SnRK2.4 and SnRK2.10 kinases found in senescing Arabidopsis leaves from plants cultivated in strictly controlled optimal conditions, that is, not subjected to any external senescence-inducing stimuli.

A transcriptome comparison between wt and *snrk2.4/2.10* showed pronounced differences in the gene expression profiles in senescent leaves only. Numerous DEGs belonging mainly to the GO categories such as response to water deprivation, response to abscisic acid, leaf senescence, and response to salt stress were down-regulated in *snrk2.4/2.10* plants. This differential expression of stress-responsive genes reflects the essence of the SnRK2.4, and −2.10 functioning in the conditioning of the plant resistance to water deficiency (McLoughlin et al., 2012; Mazur et al., 2021).

The DEGs found in our study largely coincided with the stress-related genes controlled by these kinases identified earlier (Kawa et al., 2020; Rachowka et al., 2023). As shown previously, gene expression in salinity stress and during developmental leaf senescence is controlled to some extent by the same transcription factors such as ORE1 (Balazadeh et al., 2010; Allu et al., 2014). Among the DEGs found in our study, 61 were ABA-responsive ones induced by SnRK2.4 and SnRK2.10. Abscisic acid is a phytohormone synthesized from a carotenoid precursor and the first enzyme committed specifically to ABA synthesis is 9-*cis*-epoxycarotenoid dioxygenase which cleaves an epoxycarotenoid precursor to form xanthoxin (Seo and Koshiba, 2002). At the basal level, ABA plays numerous vital roles in promoting plant growth and development, including modulation of tillering, flowering, seed development and seed maturation. ABA also confers plant tolerance to multiple stress factors (for review see Yoshida et al., 2019; Kishor et al., 2022). Endogenous and externally added ABA induces leaf senescence (Gao et al., 2016; Xu et al., 2020), which is why until now mainly the ABA-activated protein kinases have been studied as leaf senescence inducers (Matsuoka et al., 2015; Gao et al., 2016). Nevertheless, some studies have indicated the action of ABA-independent pathways in developmental leaf senescence and suggested that endogenous ABA-dependent leaf senescence requires an additional primary stress signal to be initiated (Pourtau et al., 2004; Song et al., 2016). The nature of this signal is still being investigated. Here, we show that the ABA-non-activated SnRK2.4 and SnRK2.10 act upstream of and control ABA accumulation, indicating their primary role in regulating the ABA-dependent processes in senescing leaves. Recently, Zhang et al. (2024) showed that in response to cold CaSnRK2.4, an ABA-activated SnRK2 of *Capsicum annuum*, phosphorylates the CaNAC035 transcription factor which promotes ABA accumulation by activating two ABA-biosynthesis related genes, *CaAAO3* and *CaNCED3*, in response to cold. In contrast, SnRK2.3 phosphorylates and inhibits HAT1 (HOMEODOMEIN ARABIDOPSIS THALIANA 1) (Tan et al., 2018) which represses the ABA biosynthesis genes *NCED3* and *ABA3* and allows for fine-tuning of the Arabidopsis response to drought. In senescing Arabidopsis leaves, of all the NCEDs mostly NCED2 and NCED3 redundantly participate in ABA production since the stay-green phenotype in older plants was observed only in the double *nced2/3* mutant but not in the respective single mutants (Xu et al., 2020). However, here we found that only the expression of *NCED2* was strongly (over 35-fold) reduced in leaves of *snrk2.4/2.10* plants at 45 DAS, while other ABA-biosynthesis and – degradation-related genes, including other *NCEDs (NCED3* and *NCED4),* showed only minor or no differences in expression relative to wt and most likely do not affect ABA accumulation to a large extent. This indicates that SnRK2.4 and SnRK2.10 promote ABA biosynthesis by induction of only the *NCED2* gene. The mechanism of this induction is not yet known and the identification of direct SnRK2.4 and SnRK2.10 targets, including the transcription factors involved in ABA biosynthesis, is an obvious aim of future studies.

Abscisic acid modulates multiple signaling pathways, including Mitogen-Activated Protein Kinase (MAPK) cascades which are intensively studied as being essential for diverse biotic and abiotic stress responses and maintaining proper plant development (Zhang and Zhang, 2022; Li et al., 2024). A MAPK cascade typically comprises three kinases, MAPKKK, MAPKK, and MAPK, which sequentially phosphorylate and thereby activate the downstream kinase (Ichimura et al., 2002). Of those, the MAPKKKs constitute the largest family, counting 80 members in Arabidopsis, and relatively the least known. A representative of this family, MAPKKK18, is involved in diverse ABA-regulated processes, such as seed germination, stomatal development and functioning, and drought response (Danquah et al., 2015; Mitula et al., 2015; Li et al., 2017). Notably, the complete module of the MAPKKK18/17 – MKK3 - MPK1/2/7/14 cascade has been identified. Further, MAPKKK18 has been shown to take part in both the ABA-induced and the developmentally-triggered leaf senescence (Matsuoka et al., 2015). Here we showed that MAPKKK18 is activated also in the *snrk2.4/2.10* mutant, but its activity and the level of *MAPKKK18* expression are both significantly lower in senescing leaves of the *snrk2.4/2.10* mutants than in the wt plants. Interestingly, the progression of developmental leaf senescence in a *mapkkk18* line was intermediate between those in the wt and in the *snrk2.4/2.10* mutant, suggesting that SnRK2.4 and SnRK2.10 control the MAPKKK18 cascade most likely by sustaining ABA production in senescing leaves. However, the crosstalk between the SnRK2 and MAPKKK18 signaling pathways seems more complex. First, it has been shown that the ABA-triggered activity of MAPKKK18 is at least partially mediated by an ABA-activated kinase, SnRK2.6, and inhibited by ABI1 (Abscisic acid Insensitive 1), a group A protein phosphatase type 2C (PP2C) (Mitula et al., 2015). ABI1 is a negative regulator of ABA signaling which inhibits ABA-activated SnRK2s (for review, see Fàbregas et al., 2020; Maszkowska et al., 2021) and also ABA-non-activated SnRK2s (Krzywińska et al., 2016, Yuan and Zhao, 2025).

The expression of *MAPKKK18* is induced by ABSCISIC ACID RESPONSIVE ELEMENTS-BINDING FACTORs ABF2, ABF3, and ABF4, the core ABA-signaling transcription factors. ABF4 binds directly to the *MAPKKK18* promoter (Zhao et al., 2023). The three ABFs act redundantly and the triple knock-out mutant *abf2/abf3/abf4* exhibits a stay-green phenotype after ABA treatment along with a decreased induction of *SGR1* and *SGR2* and the direct target genes *PAO1* and *NYC1* (Gao et al., 2016), which are all involved in the PAO/phyllobilin pathway of chlorophyll degradation (Zhu et al., 2017). ABF2, −3, and −4 control leaf senescence not only *via* MAPKKK18 but also independently of it (Zhao et al., 2023). ABF2, −3, and −4 are major substrates for the SnRK2.2, SnRK2.3, and SnRK2.6 kinases (Furihata et al., 2006), predominantly in the regulation of gene expression in the ABA-dependent response to osmotic stress (Yoshida et al., 2015). Also, SnRK2.4 and SnRK2.10 phosphorylate and activate ABF2 efficiently (Ruschhaupt et al., 2019; Fig. S7). These findings indicate that SnRK2.4 and SnRK2.10 could regulate ABF2–dependent gene expression directly by its phosphorylation and indirectly by promoting ABA accumulation. A schema of the postulated crosstalk among the SnRK2s signaling pathways, ABA, MAPKKK18, and of ABF-dependent and -independent gene expression in the leaf senescence is presented in Fig. 6.

**Figure 6.**
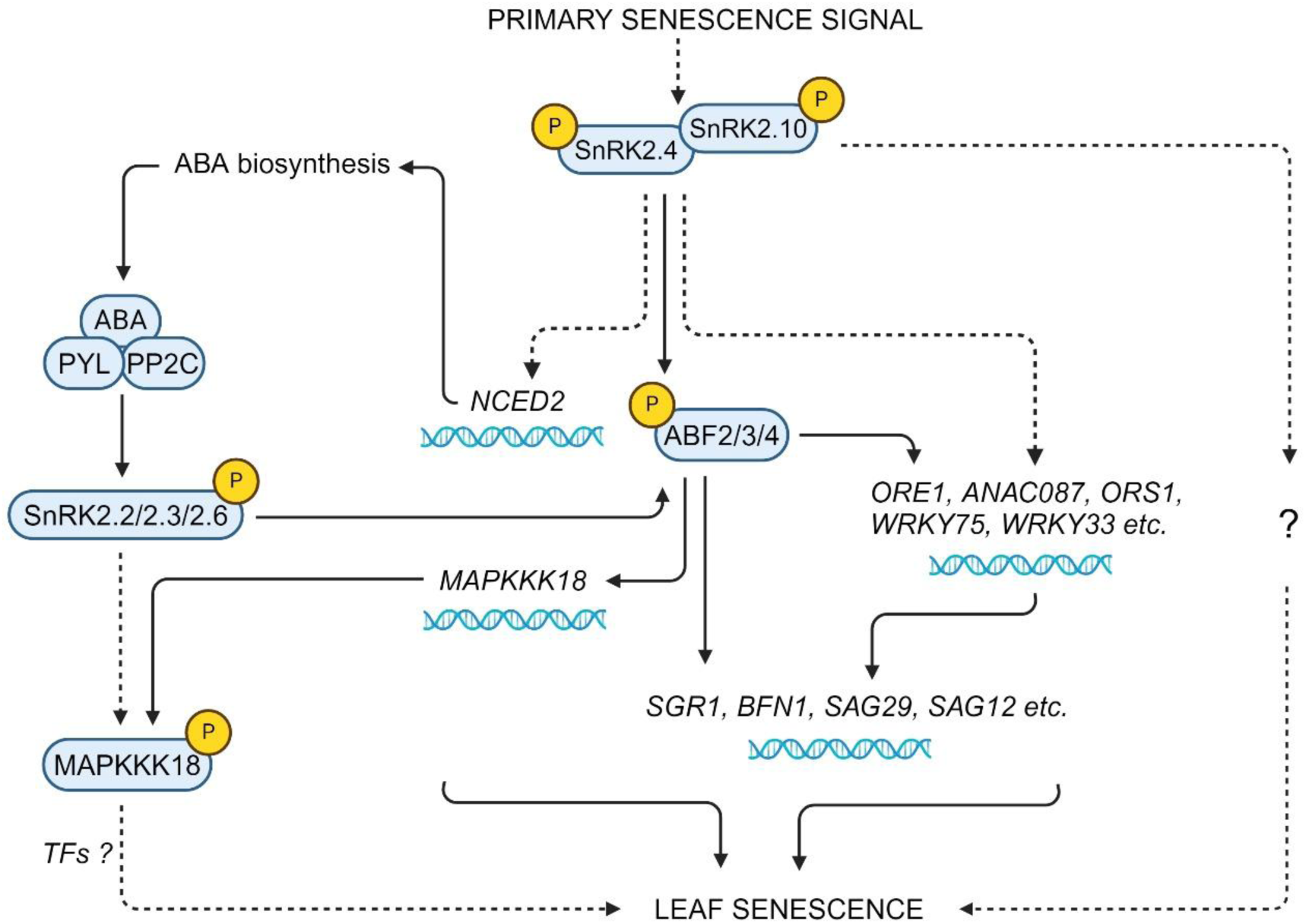
SnRK2.4- and SnRK2.10-dependent regulation of developmental leaf senescence. Schematic view of the complex crosstalk between SnRK2s signaling pathways, ABA homeostasis, MAPKKK18 activation, and regulation of ABA-dependent and -independent gene expression. The schema is based on the results published by Furihata et al. (2006), Matsuoka et al. (2015), Mitula et al. (2015), Gao et al. (2016), Zhao et al. (2023), and our current findings. Solid lines indicate well-established relations, dashed lines – putative ones.

In our study, induction of the leaf senescence by exogenously applied ABA was not affected in *snrk2.4/2.10* plants nor was the activation of ABA-induced SnRK2s, indicating that SnRK2.4 and SnRK2.10 regulate developmental leaf senescence by inducing ABA accumulation and thereby sustaining the ABA-responsive signaling pathways, but they do not affect ABA sensing or signaling as such. However, Shahzad et al. (2024) found that SnRK2.4 promoted ABA-dependent stomatal functioning, inhibition of primary root growth, and root hydraulic conductivity by phosphorylation of aquaporins, thus contributing to ABA sensing and signaling. Importantly, reconstitution of the core Arabidopsis abscisic acid signaling pathways in yeast has revealed that group 1 SnRK2s are much more sensitive to ABA than is SnRK2.6/OST1 and can respond to very low endogenous ABA concentrations (Ruschhaupt et al., 2019). Thus, SnRK2.4 could be activated in the cell even at optimal growth conditions in response to minor ABA concentration fluctuations, and stimulate ABA biosynthesis leading to the activation of multiple ABA-dependent kinases requiring high ABA concentrations for activation. Therefore, the group 1 SnRK2 kinases could function as amplifiers of the weak developmental abscisic acid signals with respect to other ABA-dependent and –independent signaling pathways.

The upstream factors and stimuli triggering developmental and age-dependent leaf senescence are relatively poorly understood. For many years it has been discussed whether sugars, the accumulation of which is strongly associated with leaf age, could function as signaling molecules to determine the onset of leaf senescence in optimal (for review see Asad et al., 2004; Kim, 2019) and stress conditions (Asad et al., 2024). The sugar deficiency occurring during dark-induced leaf senescence or following a decline in photosynthetic efficiency have been proposed to be negative regulators of leaf senescence in the middle stages of developmental leaf senescence but not in the early stages. On the other hand, a high photosynthetic activity, external supply of sugars, or blocking of sugar export from the leaf can induce leaf senescence. In the transition between leaf maturity and early developmental leaf senescence stages or abiotic stress-induced senescence, four major sugar sensors, including T6P (TREHALOSE-6-PHOSPHATE SYNTHASE 1), HXK (Hexokinase), SnRK1, and TOR (Target of Rapamycin) have been proposed to coordinate cellular carbon signaling and determine the onset of leaf senescence (for review see Asad et al., 2004; Kim, 2019). Importantly, it has been shown that in the absence of ABA SnRK2.2 and possibly also other ABA-activated SnRK2s form a SnRK1 repressor complex with PP2Cs, preventing TOR activation and growth repression under favorable conditions (Belda-Palazón et al., 2020). Further, TOR phosphorylates PYL, a component of the PYL/PYR/RCAR ABA receptor thereby disrupting the PYL association with ABA and PP2C phosphatases, and leading to the inactivation of SnRK2 kinases. In turn, the ABA-activated SnRK2s phosphorylate Raptor, a TOR-interacting regulatory protein, thereby triggering TOR complex dissociation and inhibition under stress conditions. Thus, TOR signaling represses ABA signaling and stress responses in unstressed conditions, and reciprocally ABA signaling represses TOR signaling and prevents growth under unfavorable environmental conditions (Wang et al., 2018; Belda-Palazón et al., 2020). Under drought, SnRK2.2, −2.3, and −2.6 also sustain long-distance sugar transport by phosphorylating SWEET11 and −12 (Sugars Will Eventually be Exported Transporter) proteins in an ABA-dependent manner. The phosphorylation enhances their activity, thereby increasing sucrose export from leaf cells –-the first step of phloem loading with sucrose (Chen et al., 2022). Here, we found that SnRK2.4 and SnRK2.10 are required for *SWEET2, −4, −14*, and -*15/SAG29* expression during developmental leaf senescence, which implies their involvement in sugar translocation within the cell and plant. Whether the group 1 SnRK2 kinases are involved in the phosphorylation of the sugar transporters and/or SnRK1 and TOR functioning remains unknown. The possibility of a tight association between the ABA-non-activated SnRK2s and sugar signaling/transport seems attractive and deserving of further investigation.

Summarizing, we have shown that group 1 SnRK2s control leaf development in non-stress conditions and identified the mechanisms involved. We posit that SnRK2.4 and SnRK2.10 are activated, probably at early stages of leaf senescence, by non-stress-related endogenous factors and redundantly induce *NCED2* expression causing ABA accumulation. In response to the ABA accumulation ABA-dependent signaling pathways of the MAPKKK18 module and possibly also SnRK2s from group 3 are activated and induce the expression of multiple ABA-dependent SAGs. Further, SnRK2.4, SnRK2.10, and the ABA-activated SnRK2s phosphorylate the ABF2 TF and thus control the ABF-dependent expression of *MAPKKK18* and other SAGs sustaining leaf senescence. We posit that the kinases studied relay endogenous developmental signals and/or developmental status to major ABA-dependent and –independent signaling pathways inducing stress-like responses in senescing tissues, as summarized in Fig. 6.

## MATERIALS AND METHODS

### Plant lines, growth conditions, and material collection

*Arabidopsis thaliana* T-DNA insertion lines were obtained from the Nottingham Arabidopsis Stock Centre: wt Col-0, *snrk2.10-1 (WiscDsLox233E9), snrk2.10-3 (SAIL_698_105), snrk2.4-1 (SALK_080588), snrk2.4-2 (SALK_146522), snrk2.4/snrk2.10 (snrk2.4-1/snrk2.10-1), ataf1-2 (SALK-057618),* and *mapkkk18-1 (SALK_087047).* Transgenic lines expressing chimeric SnRK2-YFP or EGFP-SnRK2 proteins, *pSnRK2.4::SnRK2.4-YFP* and *pSnRK2.10::SnRK2.10-YFP,* and *EGFP-SnRK2.4* and *EGFP-SnRK2.10* under control of a double 35S promoter were kindly provided by Prof. Christa Testerink, Wageningen University.

Plants were cultivated in a growth chamber under long-day conditions (16h light 22°C / 8h dark 21°C; light intensity approximately 110 µmol photons m^-2^s^-1^). Rosette leaves were collected at different stages of plant growth (days after sowing, DAS), immediately frozen, and stored at −80°C until analyzed. For estimation of progression in developmental leaf senescence subsequent rosette leaves were detached and photographed with a further biochemical and molecular analysis performed for the 9^th^ rosette leaves as previously described by Garapati et al. (2015). For calculation of the rosette leaf formation rate, the number of leaves in a plant was determined every five days until it ceased to increase (ca. 28 DAS). To determine flowering initiation time, the formation of the first flower bud was assessed daily in approximately 30-50 plants.

For the determination of ABA-induced kinase activity, seedlings were grown in a sterile hydroponic culture as described in Kulik et al. (2012). Ten-day-old seedlings were treated or not (control) with 100 µM ABA for up to 60 min., immediately collected, dried from the medium, frozen, and stored at −80°C until analyzed.

### ABA-induced leaf senescence assay

ABA treatment was performed as previously described in Gao et al. (2016). The third and fourth rosette leaves were detached from plants 28 DAS and floated on medium containing or not (mock) 50 or 100 µM ABA in a Petri dish for two days in continuous dim light, photographed and immediately frozen for further analysis.

### Kinase activity determination

#### Immunoprecipitation and immunocomplex kinase activity assay

The procedure was performed as described in Rachowka et al. (2023). In brief, for immunoprecipitation of SnRK2.4-YFP, SnRK2.10-YFP, or SnRK2.2/2.3/2.6 kinases, 1100 µg or 400 µg of crude protein extracted from 8^th^-10^th^ rosette leaves of plants 35 DAS or 45 DAS, respectively, was incubated with 50 µL of GFP-Trap®_A (Chromotek) resin or 2.5 µL anti-SnRK2.2/2.3/2.6 antibodies (Agrisera) bound to 25 µL of Protein A-Agarose (Santa Cruz Biotechnology) for 2.5 h, at 4°C with gentle rocking. After intensive washing, agarose beads with bound immunocomplexes were suspended in 18 µL of 20 mM Tris–HCl, pH 7.5, 150 mM NaCl supplemented with 4 µg of Myelin Basic Protein (Sigma-Aldrich) per sample. To each sample, ATP supplemented with 1 μCi of [γ^32^P]ATP in kinase buffer (25 mM Tris-HCl, pH 7.5, 5 mM EGTA, 1 mM DTT, 30 mM MgCl_2_) was added to a final concentration of 50 μM. After 40 min. of incubation at 30°C samples were mixed with 4xLaemmli sample buffer and incubated for 3 min. at 95°C with vigorous shaking. Proteins were separated on 12% SDS polyacrylamide gel and signal was detected on Medical X-ray Blue/MXBE Film (Carestream) and in a Typhoon FLA 9000 biomolecular imager (Cytiva).

The MAPKKK18 immunocomplex kinase activity assay was performed as previously described in Mitula et al. (2015) from 250 µg of crude protein extract using 1.5 µL of anti-MAPKKK18 antibody (Agrisera) per sample.

#### In-gel protein kinase activity assay

Activation of ABA-induced kinases was monitored using the in-gel kinase activity assay according to Lin et al. (2020) with ABF2 fragment, obtained as described by Furihata et al. (2006), as the kinase substrate.

The activity of EGFP-SnRK2.4 and EGFP-SnRK2.10 was determined as described in Mazur et al. (2021) using ABF2 fragment as kinase substrate.

For quantification of incorporated radioactivity the intensity of ^32^P-labeled MBP or ABF2 fragment bands was quantified using ImageJ software and normalized against Coomassie Brilliant Blue (CBB)-stained band.

### Heterologous expression and activity determination of SnRK2s

Expression in *E.coli* and purification of recombinant SnRK2s was performed according to the procedures described by Klimecka et al. (2020), and the activity assay as described by Furihata et al. (2006).

### Immunoblotting

Immunoblotting was performed using antibodies recognizing the SAG12 protein (Agrisera; AS14 2771) according to the manufacturer’s recommendation.

### RT-qPCR

Leaves were ground to a fine powder in liquid nitrogen. RNA was extracted with Trizol (Molecular Research Center) according to manufacturer’s instructions and treated with DNase I (Thermo Scientific). Reverse transcription was performed on 1 µg of RNA using the RevertAid First Strand cDNA Synthesis Kit (Thermo Scientific). The resulting cDNA was diluted 10-fold with water and 1 µL was assayed in duplicate by qPCR in a Step One Plus device (Applied Biosystems) using GoTaq® qPCR Master Mix (Promega) and specific pairs of primers (Supplemental Table S1). Transcript levels were calculated using the delta-delta Ct method and presented relative to the housekeeping gene *PEX4* (At5g25760).

### 3’ RNA sequencing

Purified RNA (500 ng) was used for reverse transcription with 50 mM barcoded oligo(dT) primers and SuperScript III polymerase. Aliquots of each reaction were pooled to create pooled libraries for 25 DAS leaves and 45 DAS leaves. The libraries were prepared according to Krzyszton et al. (2022) and Montez et al. (2024), sequenced at the Genomics Core Facility (Centre of New Technologies, University of Warsaw, Poland), and pre-analyzed as in Krzyszton et al. (2022) and Montez et al. (2024).

### Processing of 3’ RNA-seq data and differential gene expression analysis

Transcript-level counts for RNA sequencing reads were estimated by pseudoalignment with Salmon version 0.14.2 (Patro et al., 2017) using Arabidopsis TAIR10.29 transcriptome reference. Data were processed using the Three DRNAseq R package (Guo et al., 2021b). Briefly, low-expressed transcripts and genes were filtered under the rules: an expressed transcript must have ≥ 3 samples with ≥ 1 count per million reads, and an expressed gene must have at least one expressed transcript. Then, principal component analysis was used to identify whether the RNAseq data contained batch effects. The batch effect was removed using RUVr normalization (Risso et al. 2014). Read counts were normalized using Trimmed Mean of the M-values method to reduce the effect of systematic technical biases across samples. Such obtained pre-processed data were used for differential expression analysis using the limma pipeline. Differentially expressed genes (DEGs) were identified using a minimum absolute log_2_ fold change (log_2_FC) of 1 and a maximum Benjamini-Hochberg adjusted p-value of 0.05. Venn diagrams were prepared using eulerr R package (Larsson, 2018).

### Identification of hub genes

Gene networks were constructed using the STRING 11.0 platform (Szklarczyk et al. 2019). Hub genes were identified based on the calculated degree score, maximal clique centrality score, eccentricity score, edge percolated component score, or maximum neighborhood component score using cytoHubba (Chin et al. 2014) and Cytoscape 3.5 (Shannon et al. 2003). Genes identified as hub ones basing on at least two of the above scores were analyzed and clustered based on their GO annotations and plots were made using the igraph R package (Csárdi and Nepusz 2006) and Inkscape (https://inkscape.org).

### Chlorophyll a fluorescence imaging

Chlorophyll *a* fluorescence was recorded using the Maxi version of the Imaging-PAM chlorophyll fluorescence system (Heinz Walz, Germany). Detached leaves were dark-adapted for 25 min. before measurements. The fluorescence images were recorded with a resolution of 640 × 480 pixels and with the camera parameters set to avoid pixel saturation. Minimal (F_0_) fluorescence was determined using modulated blue light of 0.5 μmol photons m^-2^ s^-1^, and maximal (F_M_) fluorescence using a blue light pulse of 0.84 s and intensity of 2700 μmol photons m^-2^ s^-1^. Data were analyzed using ImagingWinGigE software and the F_V_/F_M_ parameter was calculated using the formula (F_M_ – F_0_) / F_M_.

### Quantification of chlorophyll a and b

Chlorophyll *a* and *b* content was determined according to Wellburn (1994).

### Electrolyte leakage determination

Five 9^th^ or 10^th^ rosette leaves from plants 45 DAS were immersed in tubes filled with 10 mL of miliQ water, vortexed extensively and incubated at 25°C for 30 min. The conductivity of the liquid was determined immediately after the incubation (initial conductivity) using LAQUAtwin-EC-33 conductometer (Horiba Scientific), the samples were incubated at 95°C for 15 min and cooled to 25°C, and the conductivity was determined again (total conductivity). Electrolyte leakage was expressed as (initial conductivity) : (total conductivity)x100%.

### Quantification of H_2_O_2_

Hydrogen peroxide accumulation in leaves was determined using Amplex Red reagent (Invitrogen) according to Guo et al. (2017).

### Determination of MDA

Malondialdehyde (MDA) was estimated using methods described by Janero (1990) and Hodges et al. (1999) with modifications. Leaf tissue was ground in liquid nitrogen and 100 mg of powder was mixed with 500 uL of 0.1% trichloroacetic acid (TCA) and vortexed. The samples were centrifuged for 10 min. at 10 000 x g at room temperature and 200 µL of supernatant was added to (1) 800 µL of 20% TCA / 0.01% butylated hydroxytoluene or (2) 800 µL of 20% TCA / 0.5% thiobarbituric acid. The samples were vortexed, incubated for 30 min. at 95°C and cooled to room temperature on ice. Absorbance was measured at 440, 532 and 600 nm and MDA content was calculated using the formula:

1. A = [(Abs_532 nm + TBA_ – Abs_600 nm + TBA_) - (Abs_532 nm - TBA_ – Abs_600 nm - TBA_)]
2. B = [(Abs_440 nm + TBA_ – Abs_600 nm + TBA_) x 0.0571]
3. MDA equivalents (nmol x mL^-1^) = (A-B/157) × 10^3^

### Quantification of ABA

ABA content was determined as described by Zhang et al. (2014). Briefly, 100 mg of leaves ground in liquid nitrogen was stirred overnight in darkness at 4°C with 1.5 mL of 80% methanol, 100 mg/L butylated hydroxytoluene and 0.5 g/L citric acid monohydrate. Samples were centrifuged at 1000 x g for 20 min. at 4°C and supernatant was transferred to new tubes and dried overnight under vacuum at 4°C. The dry residue was dissolved in 100 µL of methanol and 900 µL of TBS buffer (50 mM Tris pH 7.8, 0.1 mM MgCl_2_, 150 mM NaCl) and immediately analyzed. ABA was determined with a Phytodetek® Immunoassay Kit for ABA (Agdia) according to the manufacturer’s recommendations.

### Statistical analysis

Statistical analysis was performed by One-way ANOVA followed by post-hoc Tukey test (*p* < 0.05) or Student’s *t*-test in Sigma Plot 15.0 software.

The schema presented at Fig. 6 was drawn in BioRender software.

## DATA AND CODE AVAILABILITY

RNA-seq data generated in this study have been deposited at the NCBI GEO database under accession no. GSE289280.

## ACKNOWLEDGMENTS

3’ RNA sequencing was performed at the Genomics Core Facility, CeNT, University of Warsaw (RRID:SCR_022718) using a NovaSeq 6000 platform financed by the Ministry of Science and Higher Education (decision no. 6817/IA/SP/2018 of 2018-04-10). We are grateful to Prof. C. Testerink (Wageningen University, the Netherlands) for her kind gift of seeds of the SnRK2.4-GFP, SnRK2.10-GFP, *pSnRK2.4::SnRK2.4-YFP* and *pSnRK2.10::SnRK2.10-YFP* transgenic plants, to Dr. A. Ludwików (Adam Mickiewicz University, Poland) for sharing with us the *mapkkk18-1* seeds, and to A. Kasztelan, M.Sc., for help with the purification of ABF2 protein fragment by FPLC.

No conflict of interest is declared.

## AUTHOR CONTRIBUTIONS

A.K. designed the study. A.A.-M., L.P.-K., M.K., M.O., D.C., R.M., J.R., D.T., M.B., P.S., and A.K. performed experiments. A.K., S.S., and G.D. analyzed the data. A.K. wrote and all authors revised the article.

## SUPPLEMENTARY DATA

**Figure S1.**
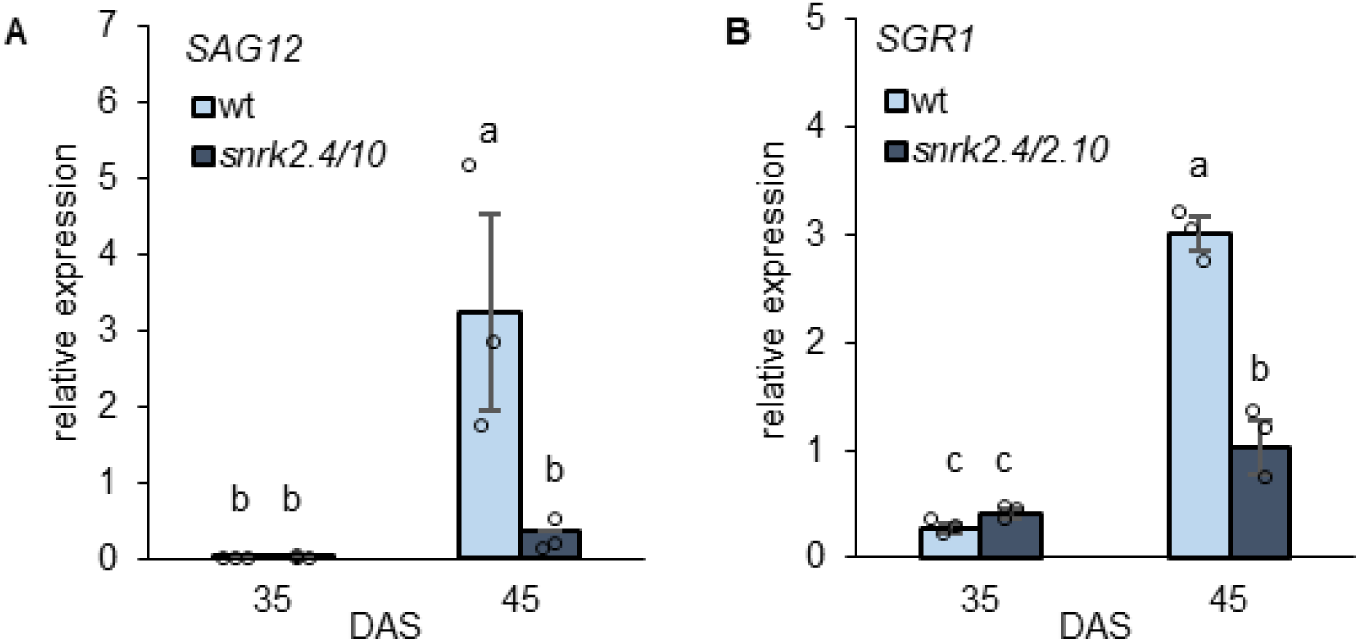
Impact of a deficit of SnRK2.4 and SnRK2.10 on the expression of developmental leaf senescence marker genes. Expression of *SAG12* (A) and *SGR1* (B) in 9^th^ rosette leaves of wt and *snrk2.4/2.10* plants 35 and 45 DAS. The transcripts were quantified by RT-qPCR relative to the *PEX4* housekeeping gene. Mean values ±SD of three independent experiments with ten leaves per sample, are shown. Different letters mark significantly different values as determined by ANOVA and Tukey post hoc test (*p* < 0.05).

**Figure S2.**
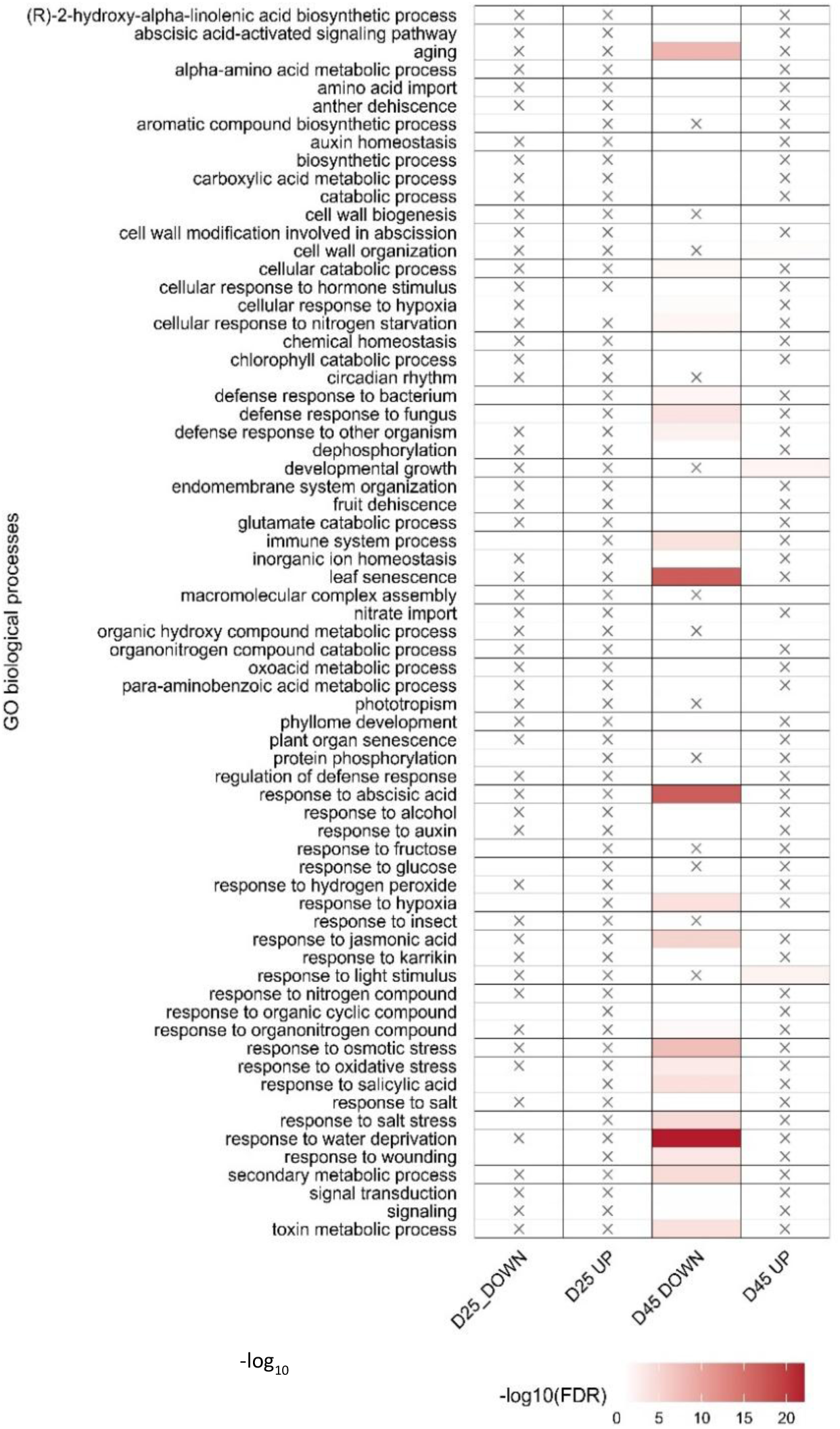
Functional analysis of DEGs in *snrk2.4/2.10* leaves. Enrichment of GO terms in “Biological process” among genes differentially expressed between *snrk2.4/2.10* and wt plants at time points indicated. Color intensity indicates the -log_10_ -transformed False Discovery Rate. All the terms shown were over-represented with a statistical significance of *p*<0.05. X means no genes detected in the term.

**Figure S3.**
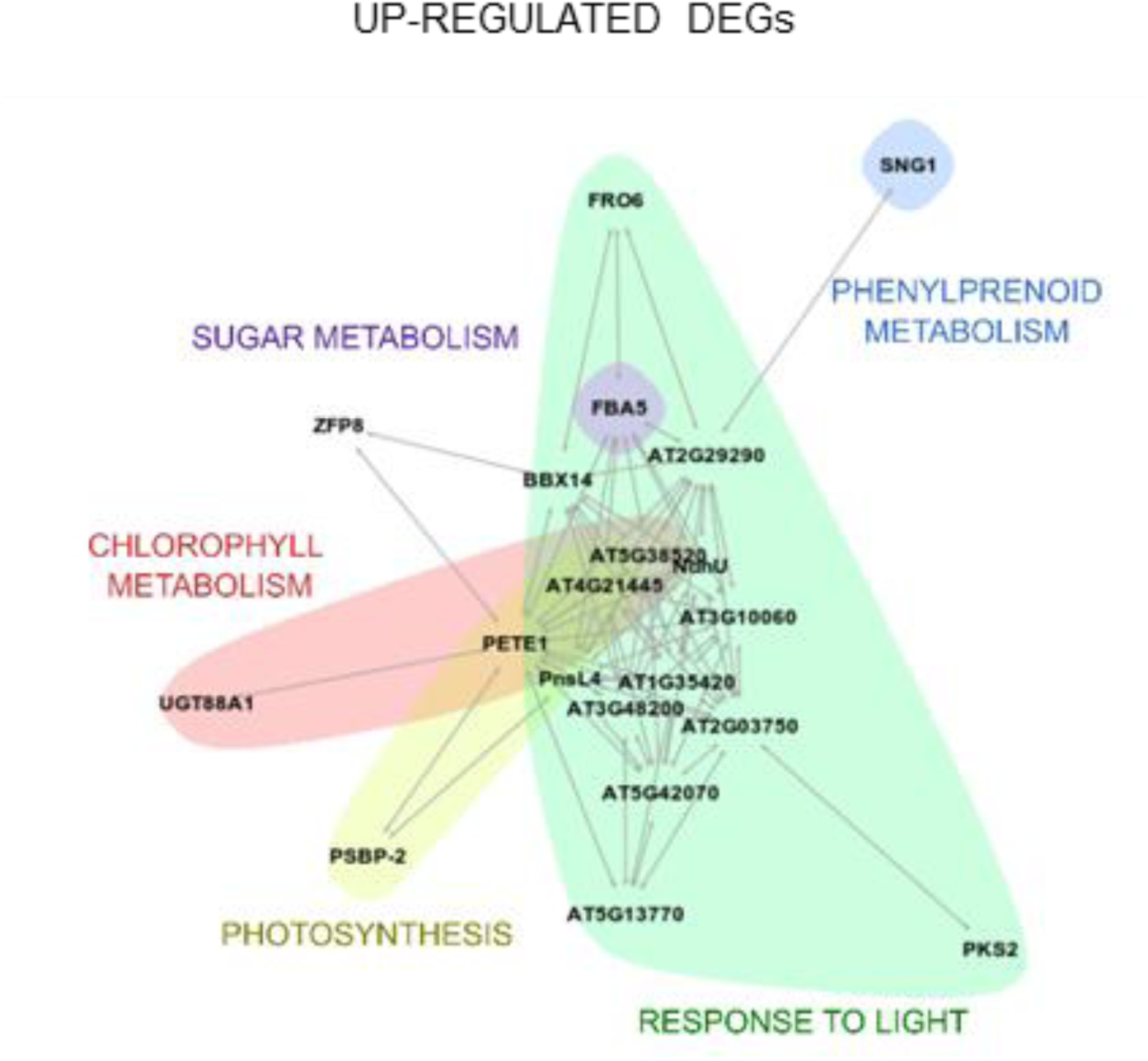
Impact of a deficit of SnRK2.4 and SnRK2.10 on the expression of hub genes. Up-regulated hub genes identified in the network analysis for *snrk2.4/2.10* 45 DAS *vs.* 25 DAS. Genes identified in wt 45 DAS *vs.* 25 DAS were subtracted from the analysis. Colors mark manually assigned groups, as labeled.

**Figure S4.**
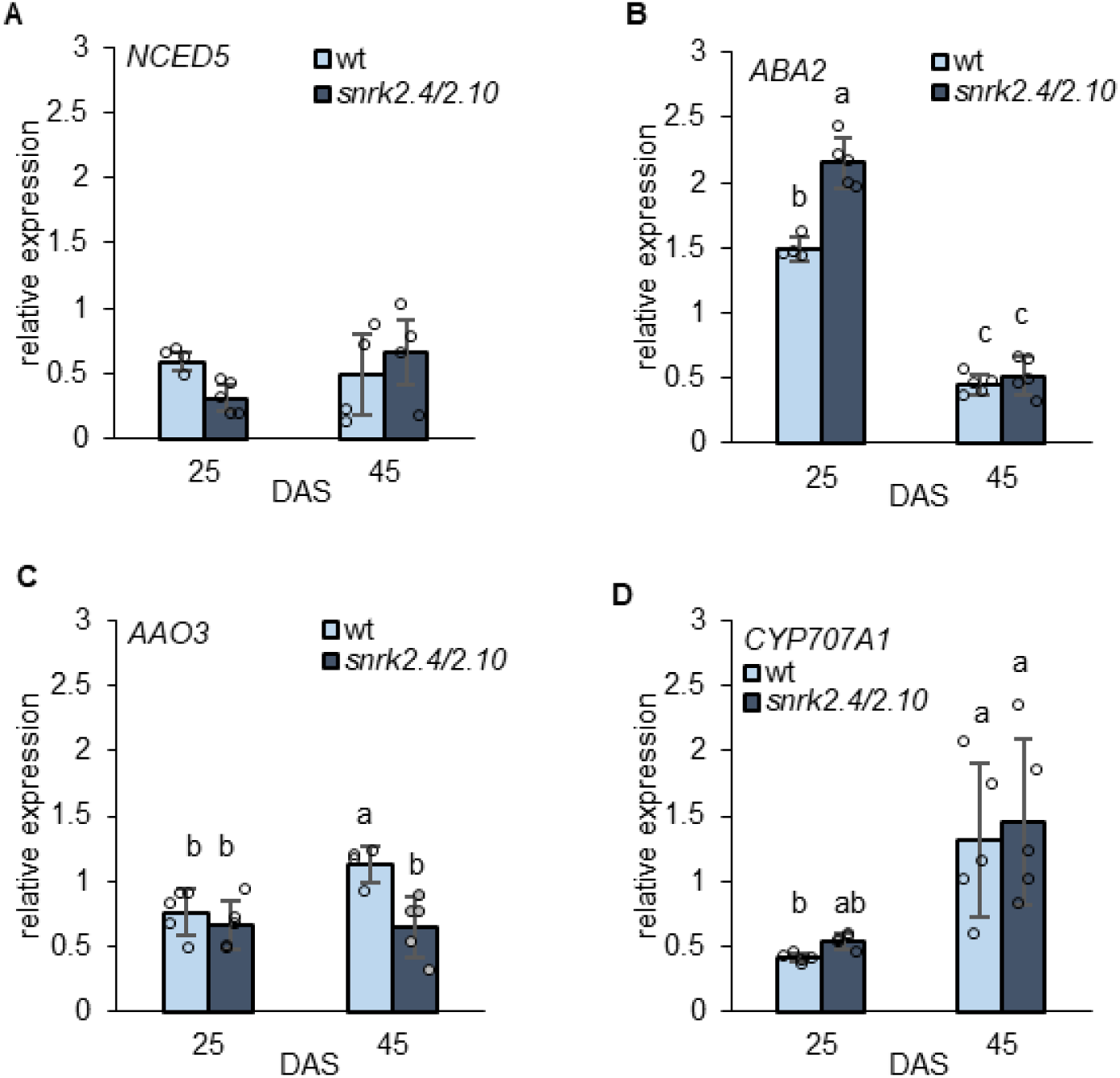
Impact of a deficit of SnRK2.4 and SnRK2.10 on the expression of ABA metabolism-related genes. Expression of *NCED5* (A), *ABA2* (B), *AAO3* (C), and *CYP707A1* (D) in 9^th^ rosette leaves of wt and *snrk2.4/2.10* plants 25 and 45 DAS. The transcripts were quantified by RT-qPCR relative to the *PEX4* housekeeping gene. Mean values ±SD of three independent experiments with ten leaves per sample, are shown. Different letters mark significantly different values as determined by ANOVA and Tukey post hoc test (*p* < 0.05).

**Figure S5.**
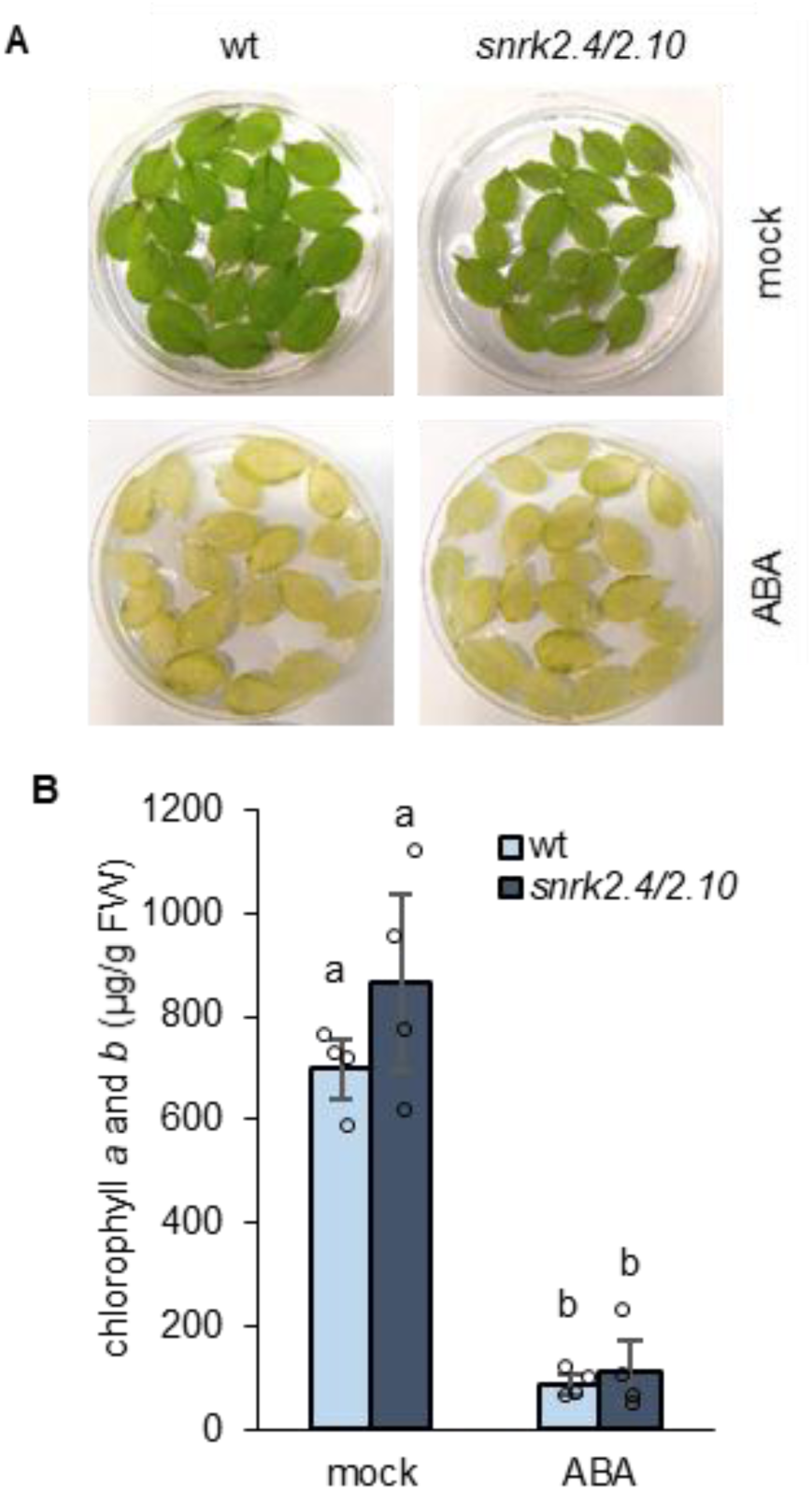
ABA-induced leaf senescence in wt and *snrk2.4/2.10* leaves. A. Phenotypes of wt and *snrk2.4/2.10* leaves after ABA treatment. Detached 3^rd^ and 4^th^ rosette leaves from plants 28 DAS were treated or not with 100 μM ABA for 2 days under continuous dim light. Representative photographs from two independent experiments performed in four replicates, with 20 leaves per sample, are shown. B. Chlorophyll *a* and *b* content in leaves of wt and *snrk2.4/2.10* plants presented in panel A. A representative graph (mean ±SD) from two independent experiments performed in four replicates with, 20 leaves per sample, is shown. Different letters mark significantly different values as determined by ANOVA and Tukey post hoc test (*p* < 0.05).

**Figure S6.**
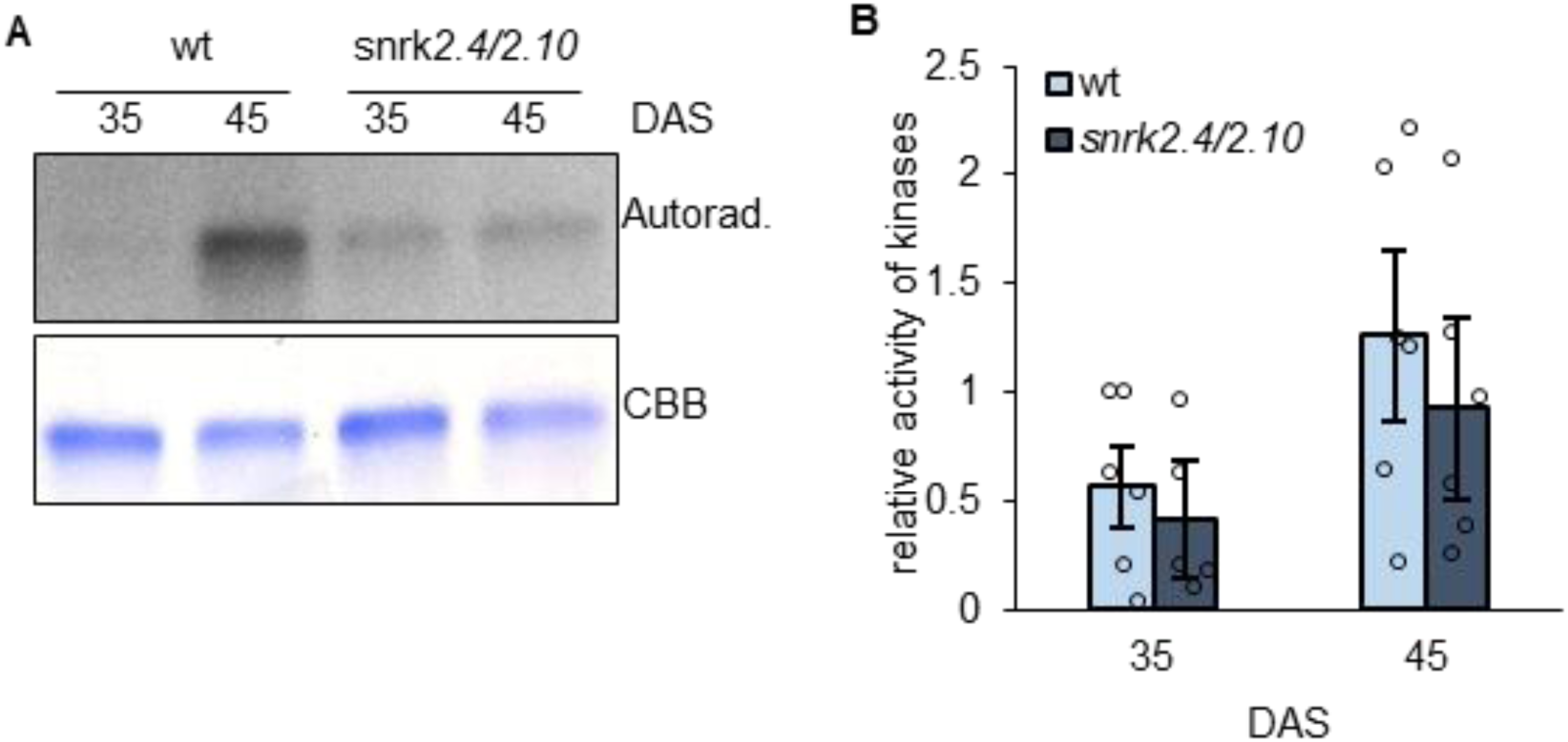
Induction of ABA-activated SnRK2s upon developmental leaf senescence. A. Activity of SnRK2.2/2.3/2.6 kinases in 8^th^-10^th^ rosette leaves of wt and *snrk2.4/2.10* plants 35 and 45 DAS. Kinase activity was determined by immunocomplex kinase activity assay using anti-SnRK2.2/2.3/2.6 antibodies and MBP as a substrate. Representative photographs from six independent experiments with eight plants per sample, are shown. Autorad. – autoradiogram. CBB – Coomassie Brilliant Blue. B. Quantification of ^32^P-labeled MBP bands phosphorylated by SnRK2.2/2.3/2.6 (shown in A). Band intensity was determined using ImageJ software and normalized against CBB-stained gel. The values shown are relative to wt plants 35 DAS. Means ±SE from six independent experiments are shown.

**Figure S7.**
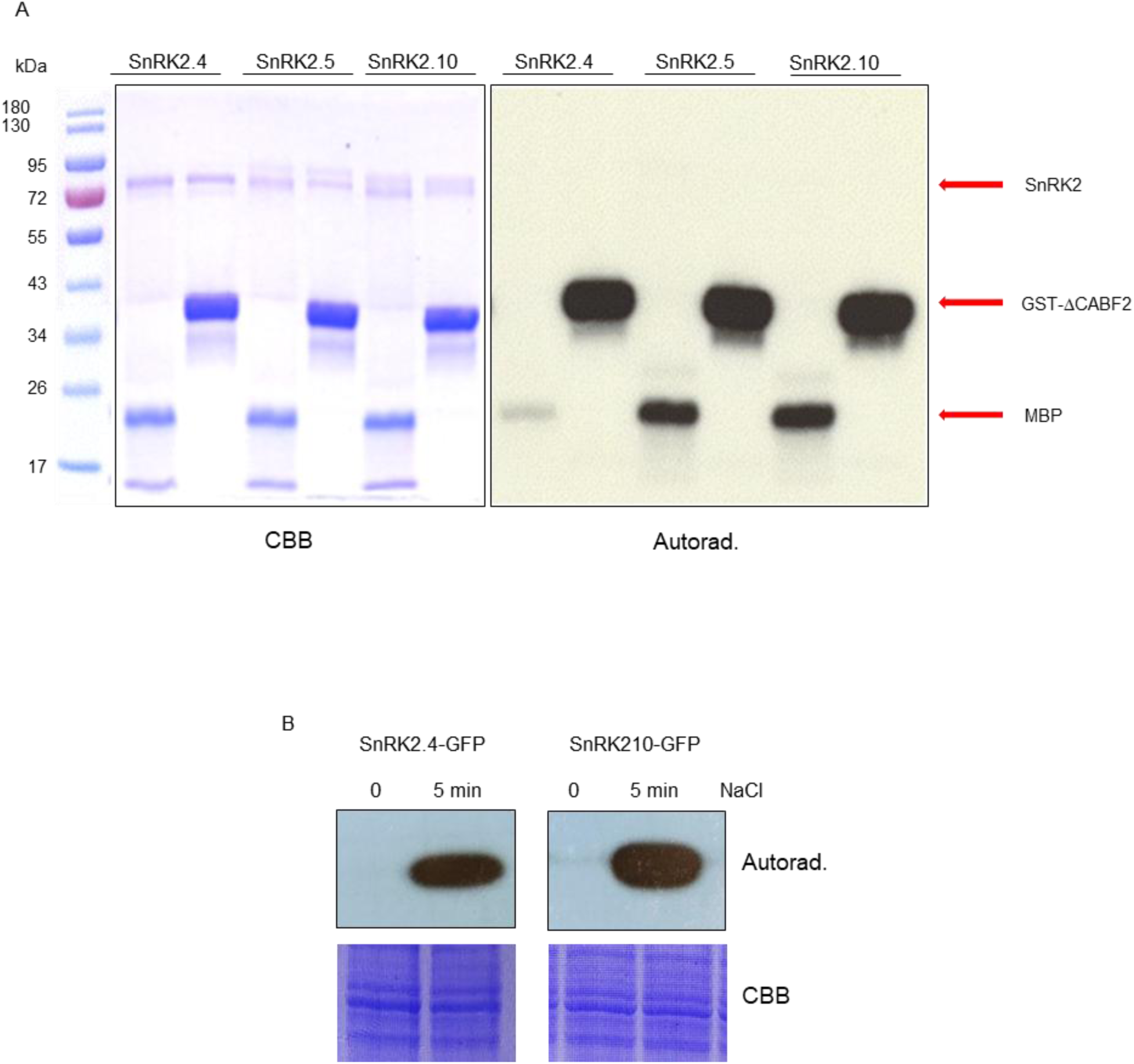
*In vitro* phosphorylation of GST-ΔCABF2 by SnRK2.4, SnRK2.5, and SnRK2.10. A. Phosphorylation of GST-ΔCABF2 (Gly73-Gln120 fragment) by recombinant SnRK2.4, SnRK2.5, and SnRK2.10 kinases. The kinases and the ΔCABF2 fragment were produced in *Escherichia coli* and used for in-solution kinase activity as assay described in Supplementary Methods. MBP was additionally used as a universal kinase substrate. Reaction products were separated by SDS-PAGE and the extent of phosphorylation was determined by autoradiography. Representative results from one of three independent experiments are shown. B. Phosphorylation of ΔCABF2 (Gly73-Gln120 fragment) by SnRK2.4-GFP or SnRK2.10-GFP obtained from plants. SnRK2.4-GFP and SnRK2.10-GFP obtained from transgenic plants before and after treatment with 250 mM NaCl for 5 min was resolved on polyacrylamide gel containing immobilized ΔCABF2 fragment. The gel was then subjected to an in-gel kinase assay. Representative results from one of three independent experiments are shown.

**Table S2.**
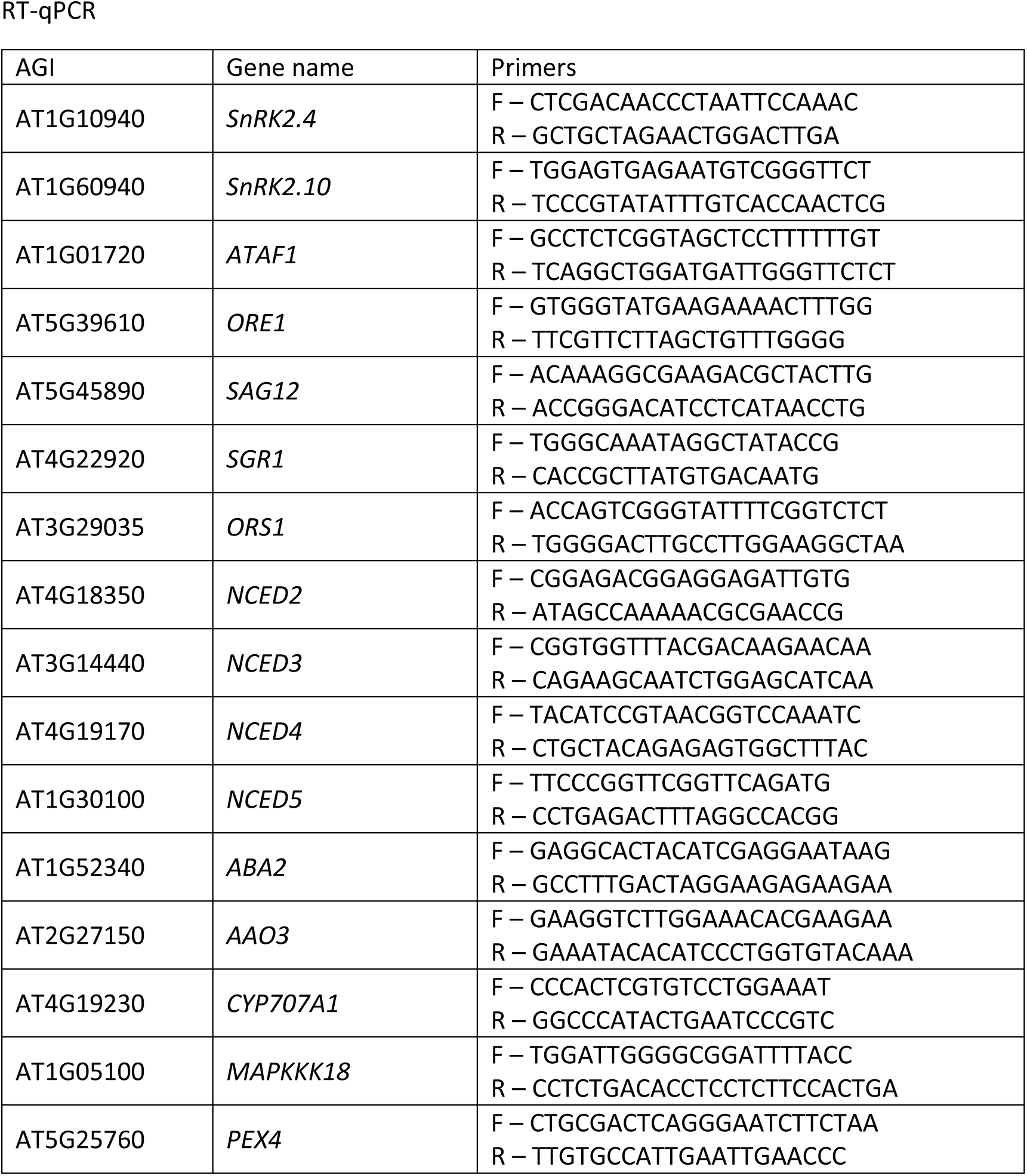
List of primers used in this study.

